# Multi-Cohort Transcriptomic Analysis Reveals Shared Inflammatory Signaling and Urothelial Dysfunction Across IC/BPS Phenotypes

**DOI:** 10.64898/2026.11.02.763762

**Authors:** Shivesh Ghura, James Griffith, Anthony J. Schaeffer, David J. Klumpp

**Affiliations:** Departments of Urology, University of Chicago Medicine and Biological Sciences, Chicago, IL USA; Department of Obstetrics and Gynecology, University of Chicago Medicine and Biological Sciences, Chicago, IL USA; Department of Psychiatry and Behavioral Neuroscience, University of Chicago Medicine and Biological Sciences, Chicago, IL USA; Microbiology-Immunology Feinberg School of Medicine, Northwestern University Chicago, IL USA

## Abstract

Interstitial cystitis/bladder pain syndrome (IC/BPS) is a heterogeneous disorder characterized by bladder pain, inflammation, and urothelial dysfunction, yet the molecular features shared across clinically defined disease phenotypes remain incompletely understood. Toll-like receptor 4 (TLR4) and acyloxyacyl hydrolase (AOAH), a regulator of TLR4 ligand activity, have been implicated in IC/BPS in clinical and preclinical mechanistic studies. Here, we performed a multi-cohort reanalysis of four independent human IC/BPS transcriptomic datasets spanning bladder tissue and urine sediment from non-ulcer/non-Hunner, ulcerative/Hunner, normal-capacity, low-capacity, non-lesion, and lesion phenotypes. We evaluated immune cell-associated expression, urothelial differentiation and barrier integrity, TLR4 and pro-inflammatory signaling, and *AOAH* expression. IC/BPS exhibited phenotype-specific immune remodeling, yet shared features were also identified. Bladder-tissue IC/BPS was consistently associated with reduced expression of urothelial differentiation and barrier markers, together with increased TLR4 recognition and pro-inflammatory signaling. *AOAH* expression was elevated across all three bladder-tissue cohorts and was associated with monocyte/macrophage and pro-inflammatory transcriptional states. TLR4 signaling similarly paralleled myeloid and inflammatory features, whereas inflammatory programs generally showed inverse relationships with urothelial barrier markers. Notably, independently defined severe phenotypes of ulcerative IC and low-capacity IC showed concordant molecular remodeling characterized by increased leukocyte and monocyte/macrophage signatures, enhanced TLR4 signaling, elevated *AOAH* expression, and reduced urothelial barrier-associated expression. Together, these findings identify reproducible molecular signatures that extend across IC/BPS phenotypes and support a model in which urothelial dysfunction and TLR4-AOAH-associated myeloid inflammatory states coexist across disease phenotypes. These results provide a molecular framework for stratifying IC/BPS beyond clinical labels and identify shared and subtype-specific molecular signatures that may inform therapeutic response evaluation and clinical management.

## INTRODUCTION

Interstitial cystitis/bladder pain syndrome (IC/BPS) is a chronic pelvic pain disorder characterized by bladder-associated pain for over eight weeks together with urinary symptoms including urinary frequency in the absence of an identifiable alternative cause [1, 2]. A persistent challenge in understanding IC/BPS is its marked clinical heterogeneity [2]. IC/BPS encompasses clinically distinct phenotypes defined by features such as Hunner lesions and/or diminished bladder capacity, yet the extent to which these phenotypes reflect distinct-versus-shared underlying biological processes remains incompletely understood [3, 4]. Transcriptomic studies of bladder tissue and urine-derived epithelial sediments have identified immune infiltration along with inflammatory and epithelial abnormalities in IC/BPS, but differences in clinical classification, specimen type, geographic population, and transcriptomic platform obscure molecular features shared across cohorts and subtypes [5–10]. Further analyses of these independently generated, clinically annotated human IC/BPS transcriptomic datasets therefore provide an opportunity to identify shared molecular programs that persist despite study heterogeneity.

Urothelial dysfunction and inflammation have each been implicated as central components of IC/BPS pathophysiology [2, 11]. The urothelium provides a specialized barrier between urine and underlying bladder tissue, and altered expression of urothelial differentiation and junctional proteins may increase exposure of sensory, stromal, and immune compartments to urinary contents [11–15]. At the same time, both innate and adaptive immune abnormalities have been reported in IC/BPS, including leukocyte infiltration and altered myeloid-, T-cell-, B-cell-, mast-cell-, and cytokine-associated responses [16–21]. Whether epithelial dysfunction and inflammation represent independent features of disease or components of a coordinated molecular state remains unresolved.

Among innate immune pathways relevant to IC/BPS, Toll-like receptor 4 (TLR4) provides a plausible connection between microbial exposure and persistent inflammation, and indeed elevated peripheral TLR4 responsiveness emerged from the NIDDK MAPP studies as a signature of UCPPS [22–25]. TLR4, together with the co-receptors CD14, LY96/MD2, and LBP, recognizes bacterial lipopolysaccharide (LPS) and initiates signaling through proximal adaptor pathways that regulate inflammatory gene expression [26]. Acyloxyacyl hydrolase (AOAH) provides an additional layer of TLR4 regulation by modifying the biological activity of LPS and thereby limiting persistent LPS– TLR4 stimulation [27]. AOAH is also strongly associated with myeloid-lineage biology, making its expression relevant both as a regulator of LPS responses and as a feature of the inflammatory cellular state [28]. Human genetic studies have additionally associated variation in AOAH with inflammatory and neurobehavioral phenotypes [29–33], whereas experimental studies identify AOAH as a genetic modifier of IC/BPS-associated symptoms [34–37]. Accordingly, rather than treating AOAH and TLR4 as a linear signaling pathway, these observations suggest the consideration of a broader TLR4-AOAH axis in which TLR4-associated inflammatory signaling and AOAH-mediated regulation may intersect within myeloid-rich disease states.

Here, we performed a harmonized reanalysis of four independent human IC/BPS transcriptomic cohorts representing diverse clinical phenotypes, specimen types, platforms, and geographic populations. We examined transcriptional features representing immune-cell states, urothelial differentiation and barrier integrity, TLR4 signaling, and *AOAH* expression to determine whether these molecular features converge across heterogeneous IC/BPS populations and clinically severe phenotypes. We further compared ulcerative and low-capacity IC, two clinically severe sub-phenotypes defined in independent cohorts, to determine whether distinct clinical classifications converge on a shared molecular state. Our analyses identify recurrent immune and inflammatory alterations accompanied by urothelial dysfunction across independent IC/BPS cohorts and position TLR4 signaling and AOAH within this broader disease-associated inflammatory state.

## RESULTS

### Four independent IC/BPS cohorts enable cross-phenotype transcriptomic comparison

We identified four publicly available human IC/BPS transcriptomic studies that included transcriptome-wide expression profiling, clinically defined IC/BPS subgroups, and an appropriate comparator population [5–10]. Three datasets were generated from studies of bladder tissue and constituted the primary tissue-based analysis, whereas the fourth dataset, GSE28242, was generated from urine sediments and provided an orthogonal specimen type (Figure 1 and Table 1). GSE11783 included healthy controls (n=6), non-ulcer IC (n=5), and ulcerative IC (n=5); GSE57560 included healthy controls (n=3), normal-capacity IC (n=9), and low-capacity IC (n=4); GSE238208 included Bacillus Calmette–Guérin (BCG)-treated comparators (n=13), and paired biopsies from Hunner lesions and non-lesion background bladder tissue obtained from patients with Hunner-type IC (n=25 patients); and GSE28242 included healthy controls (n=5), non-lesion IC (n=5), and lesion IC (n=3). These cohorts differed substantially in specimen type, transcriptomic platform, geographic origin, and clinical classification, yet permit assessment of molecular convergence across heterogeneous disease definitions. Here, each cohort was analyzed independently before cross-cohort comparison in order to preserve cohort-specific biological and technical variation.

**Figure 1.**
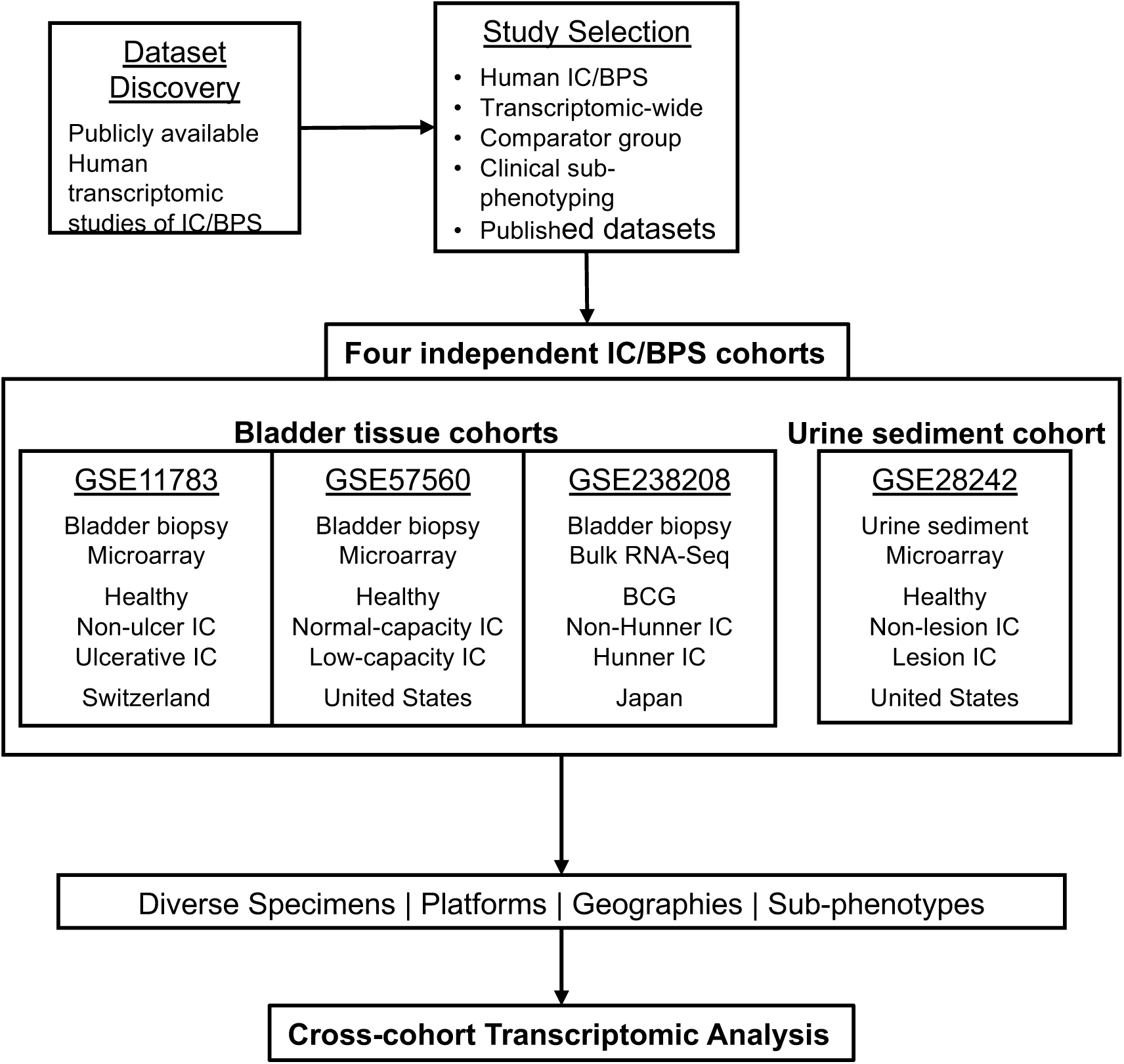
Study selection and multi-cohort analytical framework for human IC/BPS transcriptomic reanalysis. Publicly available human transcriptomic datasets of IC/BPS were evaluated for inclusion based on transcriptome-wide gene-expression profiling, availability of an appropriate comparator group, clinical sub-phenotype information, and publicly available expression data. Four independent cohorts encompassing different specimen types, transcriptomic platforms, geographic populations, and clinically defined IC/BPS sub-phenotypes were included: GSE11783, a Swiss bladder-biopsy microarray cohort comprising healthy controls (n = 6), non-ulcer IC (n = 5), and ulcerative IC (n = 5); GSE57560, a United States bladder-biopsy microarray cohort comprising healthy controls (n = 3), normal-capacity IC (n = 9), and low-capacity IC (n = 4); GSE238208, a Japanese bladder-biopsy bulk RNA-sequencing cohort comprising BCG-treated comparators (n = 13), background non-lesion biopsies (n = 25) and paired Hunner lesion biopsies (n = 25) from patients with Hunner-type IC; GSE28242, a United States urine-sediment microarray cohort comprising healthy controls (n = 5), non-lesion IC (n = 5), and lesion IC (n = 3). The three bladder-tissue cohorts formed the primary tissue-based analysis, while GSE28242 provided an orthogonal specimen type. Each cohort was analyzed independently to preserve cohort-specific biological and technical variation, followed by cross-cohort comparison of recurrent molecular features.

**Table 1.**

|  | **GEO Accession#** | **Tissue, Indication** | **Technology, Platform** | **Groups and size (n)** | **Authors, Affiliation** |
|---|---|---|---|---|---|
| 1 | **GSE11783** | Bladder biopsy, IC/BPS | Microarray, Affymetrix Human Genome U133 Plus 2.0 Array | Healthy (6)<br>Non-ulcer IC (5)<br>Ulcerative IC (5) | Gamper M, Viereck V, Geissbühler V, Eberhard J, Binder J, Moll C, Rehrauer H, Moser R<br>Functional Genomics Center, Zurich, Switzerland |
| 2 | **GSE57560** | Bladder biopsy, IC/BPS | Microarray, Agilent-039494 SurePrint G3 Human GE v2 8x60K Microarray 039381 | Healthy (3)<br>Normal capacity IC (9)<br>Low- capacity IC (4) | Colaco M, Koslov DS, Keys T, Evans RJ, Badlani GH, Erik-Andersson K, Walker SJ<br>Wake Forest University, Winston-Salem, North Carolina, USA |
| 3 | **GSE238208** | Bladder biopsy, IC/BPS | Bulk RNA-sequencing, Illumina NovaSeq 6000 | BCG (13)<br>Non-Hunner IC (25)<br>Hunner IC (25) | Akiyama Y, Luo Y, Hori M<br>The University of Tokyo, Japan |
| 4 | **GSE28242** | Urine samples, IC/BPS | Microarray, Affymetrix Human Gene 1.0 ST Array | Healthy (5)<br>Non-lesion IC (5)<br>Lesion IC (3) | Blalock EM, Korrect GS, Stromberg AJ, Erickson DR<br>University of Kentucky, KY, USA |

To characterize shared and phenotype-specific molecular features, we defined transcriptional module signatures on the basis of known markers representing general and basal/regenerative urothelium, smooth muscle, fibroblast/stroma, endothelium, pan-leukocytes, monocytes/macrophages, neutrophils, T cells, B cells, and mast cells (Table 2). We additionally evaluated a urothelial barrier module, TLR4 receptor/co-receptor recognition and proximal signaling modules, pro-inflammatory and regulatory signaling modules, *AOAH* expression, and calprotectin genes *S100A8* and *S100A9*, which encode the inflammatory proteins comprising calprotectin. Within each cohort, gene expression was standardized across samples using gene-wise z-scores, and module scores were calculated as the mean standardized expression of available genes within each signature. All analyses were performed independently within each cohort to account for differences in specimen types and transcriptomic platforms.

**Table 2.**

| **Category** | **Feature / Module** | **Molecular Markers** |
|---|---|---|
| **Cell type** | General urothelium | EPCAM, KRT7, KRT8, KRT18, KRT19 |
|  | Basal/regenerative urothelium | KRT5, KRT14, KRT17, TP63 |
|  | Smooth muscle | ACTA2, TAGLN, MYH11, CNN1, DES |
|  | Fibroblast/stroma | COL1A1, COL1A2, COL3A1, DCN, LUM, COL6A1 |
|  | Endothelium | PECAM1, VWF, KDR, EMCN, ENG |
|  | Pan-leukocyte | PTPRC, CORO1A, LST1, TYROBP, FCER1G |
|  | Monocyte/macrophage | LST1, TYROBP, FCER1G, AIF1, CD68, CTSS |
|  | Neutrophil | FCGR3B, CSF3R, CEACAM8, MPO, ELANE, CTSG |
|  | T cell | CD3D, CD3E, CD3G, TRAC, TRBC1 |
|  | B cell | CD79A, CD79B, MS4A1, CD37, CD74 |
|  | Mast cell | TPSAB1, TPSB2, CPA3, KIT, MS4A2, HDC |
| **Urothelial integrity** | Urothelial differentiation | UPK1A, UPK1B, UPK2, UPK3A, KRT20 |
|  | Tight junction/barrier | TJP1, OCLN, CLDN4, CLDN8 |
| **TLR4 signaling** | TLR4 receptor/co-receptor recognition | TLR4, CD14, LY96, LBP |
|  | TLR4 proximal signaling module | TIRAP, MYD88, TICAM2 / TRAM, TICAM1 / TRIF |
|  | Pro-inflammatory module | IL1B, IL6, TNF, CCL2, CXCL8, PTGS2, MMP9 |
|  | Regulatory module | IL4, IL4R, IL10, IL10RA, STAT6, SOCS3, TGFB1 |
| **AOAH** | AOAH | AOAH |
| **Calprotectin** | Calprotectin | S100A8, S100A9 |

### IC/BPS phenotypes exhibit heterogeneous immune remodeling with recurrent myeloid involvement

Across four independent cohorts, IC/BPS was associated with heterogeneous changes in cell type-associated transcriptional signatures, encompassing urothelial, stromal, and immune compartments (Figure 2A). The heatmap summarizes the direction and magnitude of disease-associated differences relative to the corresponding comparator group within each cohort, with red indicating increased and blue indicating decreased signature scores. For example, in GSE11783, both non-ulcer and ulcerative IC exhibited increased pan-leukocyte, monocyte/macrophage, T-cell, and B-cell signatures relative to healthy controls, whereas general and basal urothelial signatures were reduced, particularly in ulcerative IC (Supplementary Figure 1). Similarly, GSE57560 revealed increased immune-associated signatures in low-capacity IC, including monocyte/macrophage, T-cell, and B-cell features, accompanied by reduced general and basal urothelial signatures (Supplementary Figure 2).

**Figure 2.**
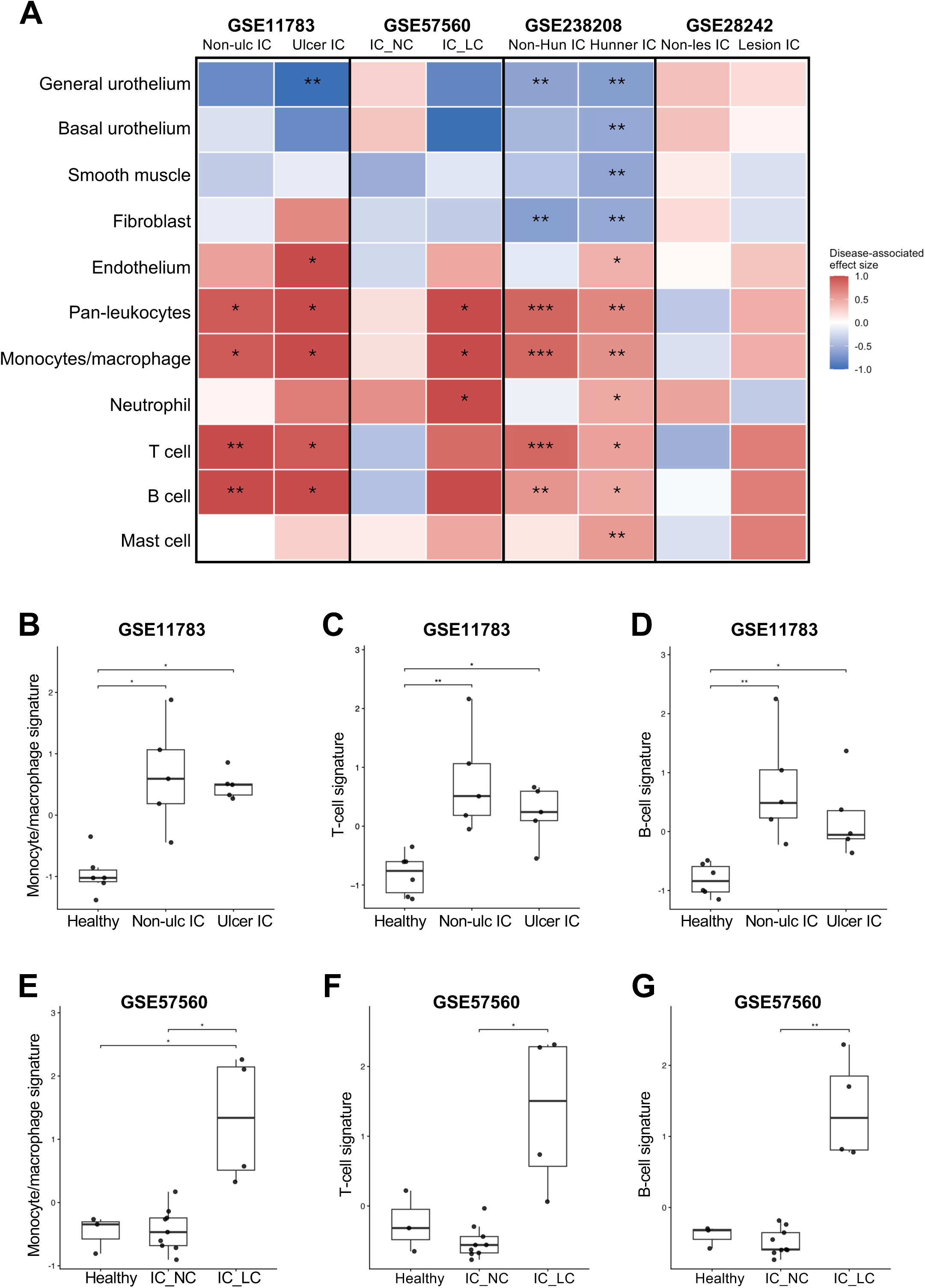
IC/BPS sub-phenotypes exhibit heterogeneous immune remodeling with shared myeloid involvement. (A) Disease-associated changes in cell type–associated transcriptional signatures across four independent IC/BPS cohorts. Signature scores were calculated using predefined marker sets representing general and basal/regenerative urothelium, smooth muscle, fibroblast/stroma, endothelium, pan-leukocytes, monocytes/macrophages, neutrophils, T cells, B cells, and mast cells (Table 2). Each IC/BPS sub-phenotype was compared with its cohort-specific comparator: healthy controls for GSE11783, GSE57560, and GSE28242, and Bacillus Calmette–Guérin (BCG)-treated comparators for GSE238208. Heatmap colors represent rank-biserial effect sizes derived from pairwise Wilcoxon rank-sum comparisons, with positive values indicating higher and negative values indicating lower signature scores in IC/BPS relative to the corresponding comparator. Sample-level distributions for all 11 signatures across the four cohorts are presented in Supplementary Figures 1–4. (B–D) Monocyte/macrophage (B), T-cell (C), and B-cell (D) signature scores in GSE11783 (healthy, n = 6; non-ulcer IC, n = 5; ulcerative IC, n = 5). (E– G) Corresponding monocyte/macrophage (E), T-cell (F), and B-cell (G) signature scores in GSE57560 (healthy, n = 3; normal-capacity IC, n = 9; low-capacity IC, n = 4). Individual points represent biological samples; boxes indicate the median and interquartile range, with whiskers extending to 1.5 × the interquartile range. Overall group differences were assessed using Kruskal– Wallis tests, followed by Dunn’s post hoc pairwise comparisons with Benjamini–Hochberg correction for multiple testing. Only significant pairwise comparisons are displayed.

These patterns were not uniform across datasets. In GSE238208, both background non-lesion and Hunner lesion biopsies exhibited increased leukocyte and myeloid-associated signatures relative to BCG-treated comparators (Supplementary Figure 3). In contrast, GSE28242 showed a distinct pattern in urine sediments, with increased T- and B-cell associated signatures in lesion-associated IC/BPS but less consistent changes across other cellular compartments (Supplementary Figure 4). Together, these findings illustrate the heterogeneous tissue-associated transcriptional remodeling across IC/BPS phenotypes, with recurrent immune enrichment and variable changes in urothelial and stromal compartments (Supplementary Figures 1–4).

Individual cell type-associated transcriptional scores further illustrate these differences (Figure 2B– G). In GSE11783, monocyte/macrophage scores were increased in both non-ulcer and ulcerative IC relative to healthy controls (Figure 2B). T-cell and B-cell signatures were similarly elevated, with particularly prominent increases in non-ulcer IC (Figures 2C, D). In GSE57560, low-capacity IC exhibited markedly increased monocyte/macrophage scores compared with healthy and normal-capacity groups (Figure 2E), accompanied by increased T-cell and B-cell signatures relative to normal-capacity IC (Figures 2F, G). Thus, although the magnitude and distribution of immune-associated transcription varied across clinical phenotypes, monocyte/macrophage-associated expression emerged as a recurrent feature of IC/BPS across independent cohorts.

### Urothelial barrier-associated expression is reduced across bladder-tissue IC/BPS cohorts

We next examined whether clinically distinct IC/BPS groups shared evidence of urothelial dysfunction. A predefined nine-gene urothelial barrier module combined differentiation-associated genes (*UPK1A, UPK1B, UPK2, UPK3A,* and *KRT20*) with tight-junction/barrier genes (*TJP1, OCLN, CLDN4,* and *CLDN8*). In GSE11783, urothelial barrier scores were reduced in both non-ulcer and ulcerative IC relative to healthy controls, with the lowest scores observed in ulcerative IC (Figure 3A). Gene-level analysis revealed marked reductions in uroplakin genes and *KRT20*, alongside decreases in selected tight junction genes, including *OCLN* and *CLDN4* (Supplementary Figure 5).

**Figure 3.**
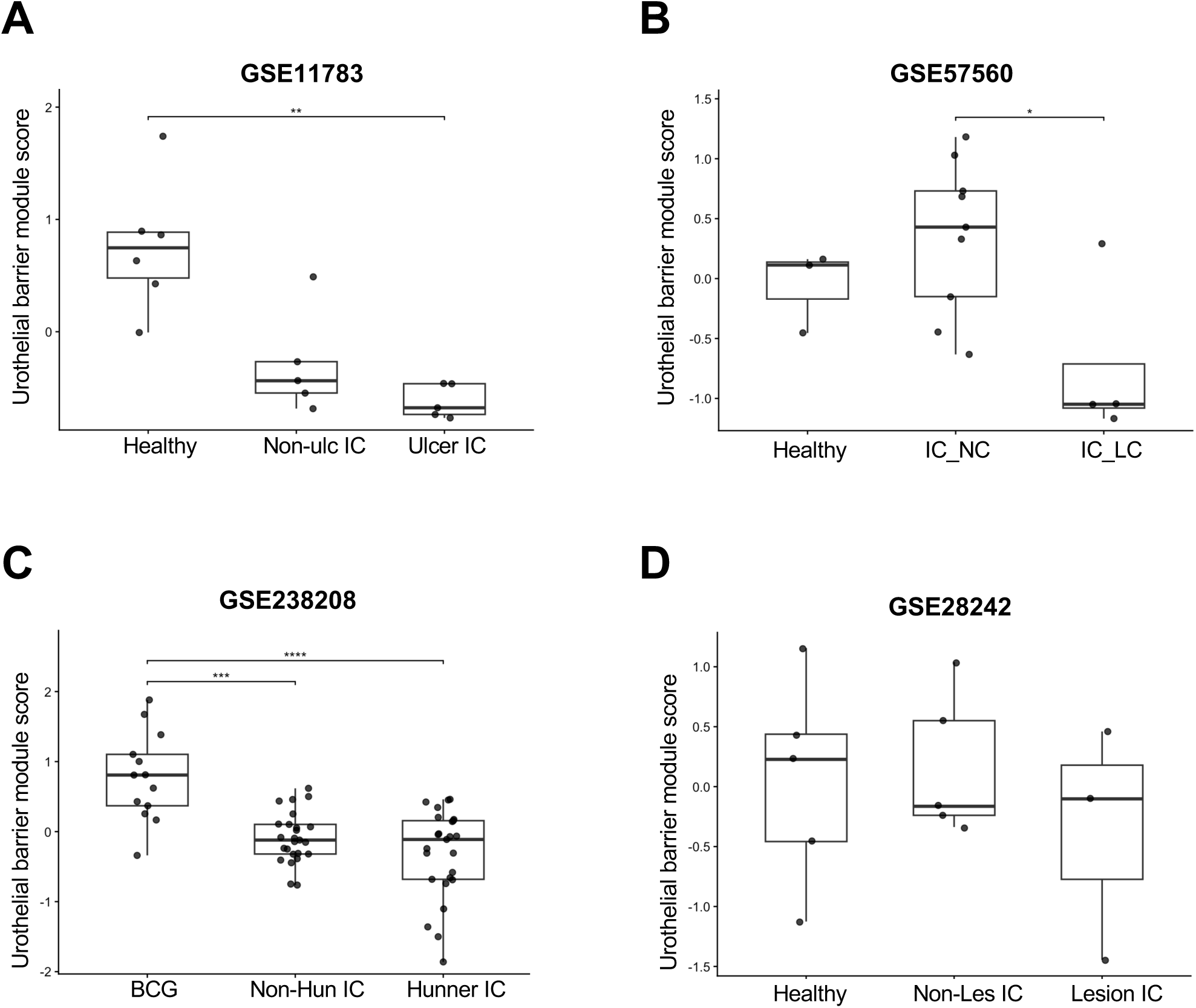
Urothelial barrier-associated expression is reduced across bladder-tissue IC/BPS cohorts. Urothelial barrier integrity was evaluated using a predefined nine-gene module comprising urothelial differentiation markers (UPK1A, UPK1B, UPK2, UPK3A, KRT20) and tight-junction/barrier markers (TJP1, OCLN, CLDN4, CLDN8). Module scores were calculated independently within each cohort from standardized gene expression; expression of the individual genes comprising the module is shown in Supplementary Figures 5-8. (A) GSE11783 bladder-biopsy cohort: healthy controls (n=6), non-ulcer IC (n=5), and ulcerative IC (n=5). (B) GSE57560 bladder-biopsy cohort: healthy controls (n=3), normal-capacity IC (n=9), and low-capacity IC (n=4). (C) GSE238208 bladder-biopsy cohort: BCG-treated comparators (n=13), background non-lesion biopsies (n = 25) and paired Hunner lesion biopsies (n = 25) from patients with Hunner-type IC. (D) GSE28242 urine-sediment cohort: healthy controls (n=5), non-lesion IC (n=5), and lesion IC (n=3). Reduced urothelial barrier scores were observed in IC/BPS groups across the three bladder-tissue cohorts (A–C), whereas significant differences were not detected in the urine-sediment cohort (D). Individual points represent biological samples; boxes indicate the median and interquartile range, with whiskers extending to 1.5 × the interquartile range. Overall group differences were assessed using Kruskal–Wallis tests followed by Dunn’s post hoc pairwise comparisons with Benjamini–Hochberg correction for multiple testing. Only significant pairwise comparisons are shown. *adjusted p < 0.05, **adjusted p < 0.01, ***adjusted p < 0.001.

Similarly, in GSE57560, low-capacity IC exhibited reduced barrier-associated expression compared with normal-capacity IC (Figure 3B), with lower expression of several urothelial differentiation markers and more variable changes among tight-junction genes (Supplementary Figure 6). In GSE238208, both background non-lesion and Hunner lesion biopsies exhibited significantly reduced barrier scores relative to BCG-treated comparators (Figure 3C). These changes were accompanied by reduced expression of multiple uroplakin genes and *KRT20*, whereas tight junction genes showed less consistent changes (Supplementary Figure 7).

In contrast to datasets derived from bladder tissue, significant differences in the composite barrier module were not detected in the urine-sediment cohort GSE28242 (Figure 3D), where individual barrier-associated genes also showed heterogeneous expression patterns (Supplementary Figure 8). This distinction is consistent with the different cellular composition and sampling characteristics of urine sediments relative to intact bladder tissue and supports use of the urine cohort as an orthogonal rather than directly equivalent specimen. Collectively, these results identify reduced urothelial differentiation and barrier-associated transcription as a reproducible feature of IC/BPS bladder tissue.

### TLR4 recognition and pro-inflammatory signaling are increased across IC/BPS phenotypes

Because innate inflammatory responses have been implicated in IC/BPS previously [23–25], we next evaluated transcriptional modules representing TLR4 receptor/co-receptor recognition, proximal TLR4 signaling, downstream pro-inflammatory signaling, and regulatory signaling. The TLR4 recognition module comprised *TLR4, CD14, LY96,* and *LBP*, while the pro-inflammatory module included *IL1B, IL6, TNF, CCL2, CXCL8, PTGS2,* and *MMP9*. TLR4 recognition scores were increased across the three bladder-tissue cohorts, with particularly prominent increases in ulcerative IC (GSE11783), low-capacity IC (GSE57560), and Hunner lesion biopsies (GSE238208) (Figure 4A–C). Pro-inflammatory module scores showed a similar pattern, with elevated expression in ulcerative IC and low-capacity IC relative to their respective controls (Figure 4D,E). In GSE238208, pro-inflammatory scores were more variable, with no significant differences detected among the groups (Figure 4F). In the urine-sediment cohort GSE28242, TLR4 recognition and pro-inflammatory scores were generally higher in lesion IC, although considerable within-group variability was observed (Supplementary Figure 9). TLR4 proximal signaling showed less consistent changes across cohorts, with significant increases in both background non-lesion and Hunner lesion biopsies relative to BCG-treated comparators in GSE238208 (Supplementary Figure 10). Regulatory module scores also varied across cohorts, with higher scores observed in ulcerative IC and low-capacity IC, but no consistent pattern of statistical differences (Supplementary Figure 11).

**Figure 4.**
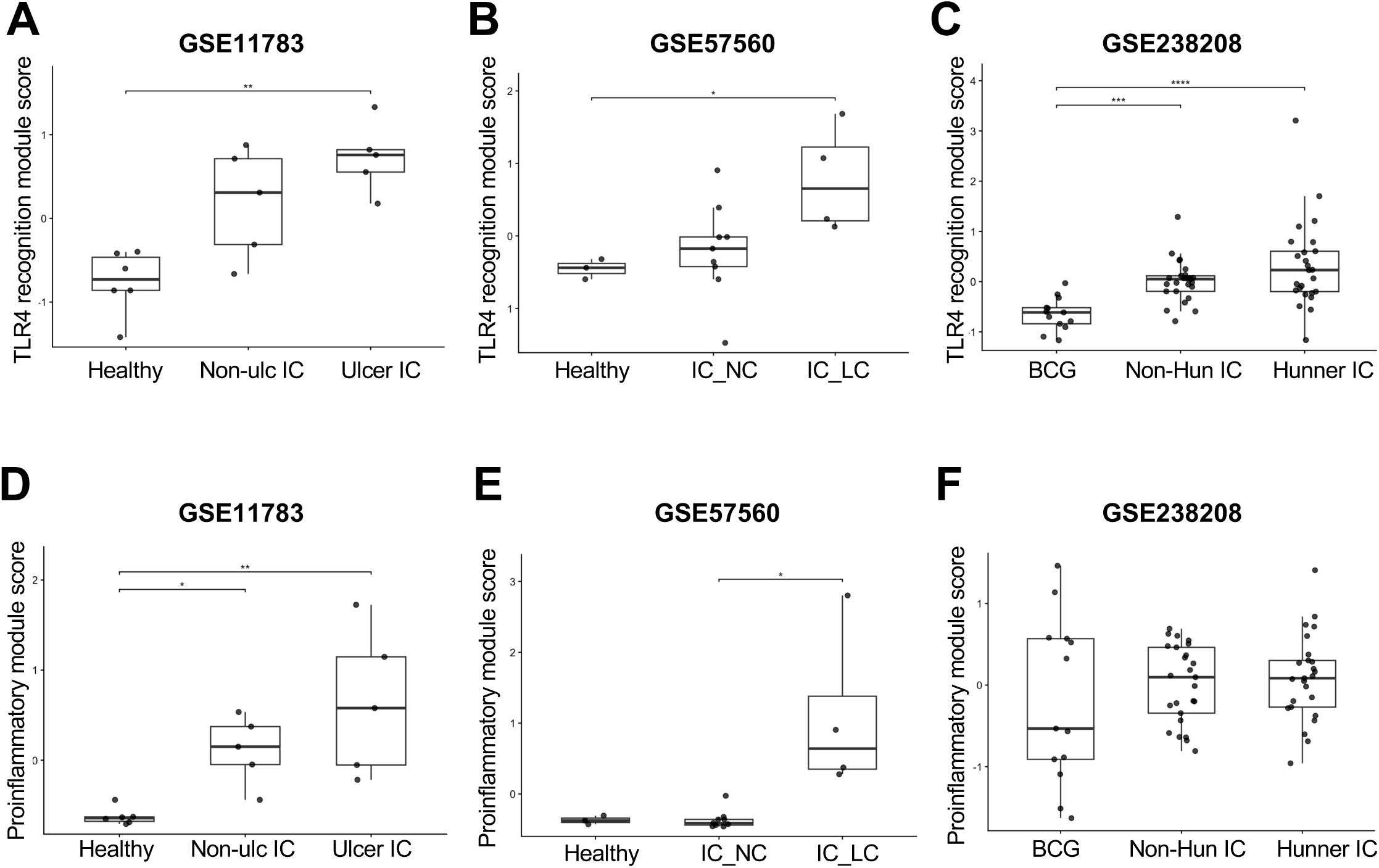
TLR4 recognition and pro-inflammatory signaling are increased across IC/BPS sub-phenotypes. TLR4-associated signaling was evaluated using predefined transcriptional modules representing receptor/co-receptor recognition (*TLR4, CD14, LY96, LBP*), proximal TLR4 signaling (*TIRAP, MYD88, TICAM1*), pro-inflammatory signaling (*IL1B, IL6, TNF, CCL2, CXCL8, PTGS2, MMP9*), and regulatory signaling (*IL4, IL4R, IL10, IL10RA, STAT6, SOCS3, TGFB1*) (Table 2). (A–C) TLR4 receptor/co-receptor recognition module scores in GSE11783 (A; healthy, n = 6; non-ulcer IC, n = 5; ulcerative IC, n = 5), GSE57560 (B; healthy, n = 3; normal-capacity IC, n = 9; low-capacity IC, n = 4), and GSE238208 (C; BCG-treated comparators, n = 13; background non-lesion biopsies (n = 25) and paired Hunner lesion biopsies (n = 25) from patients with Hunner-type IC). (D–F) Pro-inflammatory module scores in the corresponding GSE11783 (D), GSE57560 (E), and GSE238208 (F) cohorts. Corresponding TLR4 recognition and pro-inflammatory analyses for the urine-sediment cohort, GSE28242, are shown in Supplementary Figure 9. TLR4 proximal-signaling and regulatory-module analyses for all four cohorts are shown in Supplementary Figures 10 and 11, respectively. Module scores were calculated independently within each cohort from standardized expression of the constituent genes. Individual points represent biological samples; boxes indicate the median and interquartile range, with whiskers extending to 1.5 × the interquartile range. Overall group differences were assessed using Kruskal–Wallis tests followed by Dunn’s post hoc pairwise comparisons with Benjamini–Hochberg correction for multiple testing. Only significant pairwise comparisons are shown. *adjusted p < 0.05, **adjusted p < 0.01, ***adjusted p < 0.001.

Given the potential contribution of endogenous inflammatory mediators to TLR4-associated signaling, we also examined a calprotectin-associated transcriptional module comprising *S100A8* and *S100A9*. Calprotectin module scores were elevated in both non-ulcer and ulcerative IC relative to healthy controls in GSE11783 and in Hunner lesion biopsies relative to BCG-treated comparators in GSE238208 (Supplementary Figure 13A,C). In contrast, GSE57560 showed little variation in calprotectin expression across clinical groups, while GSE28242 exhibited greater within-group variability without clear phenotype-associated differences (Supplementary Figure 13B, D). Together, these findings identify increased TLR4 recognition-associated and pro-inflammatory gene expression across multiple IC/BPS phenotypes, with variation in the magnitude and statistical significance of these changes among cohorts.

### *AOAH* expression is increased across independent bladder-tissue IC/BPS cohorts

We next examined *AOAH*, which encodes an LPS-deacylating enzyme that modulates the duration and magnitude of LPS–TLR4 signaling and is strongly associated with myeloid cell biology. *AOAH* expression was increased in IC/BPS relative to cohort-specific comparator groups in each of the three primary bladder-tissue cohorts (Figure 5). In GSE11783, increased *AOAH* expression was observed in both non-ulcer and ulcerative IC relative to healthy controls (Figure 5A). In GSE57560, *AOAH* was particularly elevated in low-capacity disease (Figure 5B). In GSE238208, both Hunner lesion and background non-lesion tissue showed increased expression relative to the BCG-treated comparators (Figure 5C). In the urine-sediment cohort GSE28242, *AOAH* expression was highest in lesion IC and lowest in non-lesion IC, with variability among healthy controls (Supplementary Figure 12). Together, these findings identify increased *AOAH* expression as a recurrent feature across independent bladder-tissue IC/BPS cohorts.

**Figure 5.**
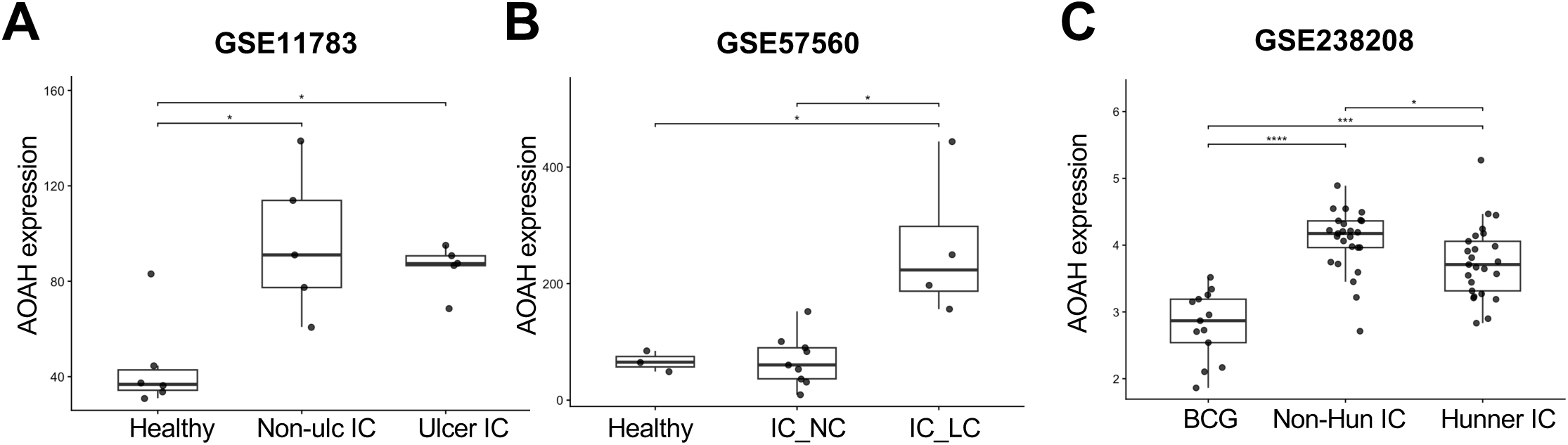
AOAH expression is elevated across bladder-tissue IC/BPS cohorts and clinically distinct sub-phenotypes. AOAH expression was evaluated independently across the three primary bladder-tissue cohorts. (A) GSE11783: healthy controls (n=6), non-ulcer IC (n=5), and ulcerative IC (n=5). (B) GSE57560: healthy controls (n=3), normal-capacity IC (n=9), and low-capacity IC (n=4). (C) GSE238208: BCG-treated comparators (n=13), background non-lesion biopsies (n = 25) and paired Hunner lesion biopsies (n = 25) from patients with Hunner-type IC. AOAH expression was increased in IC/BPS relative to the respective comparator groups across all three bladder-tissue cohorts, including independently defined ulcerative/Hunner and low-capacity disease phenotypes. AOAH expression in the orthogonal GSE28242 urine-sediment cohort is shown in Supplementary Figure 12. Individual points represent biological samples; boxes indicate the median and interquartile range, with whiskers extending to 1.5 × the interquartile range. Overall group differences were assessed using Kruskal–Wallis tests followed by Dunn’s post hoc pairwise comparisons with Benjamini–Hochberg correction for multiple testing. Only significant pairwise comparisons are shown. *adjusted p < 0.05, **adjusted p < 0.01, ***adjusted p < 0.001.

### AOAH and TLR4-associated features track inflammatory and myeloid transcriptional states

To examine relationships among *AOAH*, TLR4 signaling, and broader disease-associated transcriptional changes, we calculated within-cohort Spearman rank correlations between *AOAH* expression and predefined molecular module scores, as well as among the module scores themselves (Figure 6A). *AOAH* expression correlated positively with the pro-inflammatory module in all four datasets, while TLR4 recognition scores showed similarly positive associations with pro-inflammatory module. In contrast, *AOAH*, TLR4 recognition, and pro-inflammatory scores generally exhibited inverse relationships with urothelial barrier-associated expression, although the strength of these associations varied across cohorts (Figure 6A). Expanded analyses incorporating TLR4 proximal and regulatory signaling demonstrated additional cohort-dependent associations (Supplementary Figure 14). Calprotectin-associated transcription showed modest positive correlations with *AOAH* and pro-inflammatory signaling in several cohorts, whereas its relationship with urothelial barrier expression was less consistent (Supplementary Figure 15). Sample-level scatterplots further illustrated the direction and variability of these relationships across clinical subgroups (Supplementary Figure 16).

**Figure 6.**
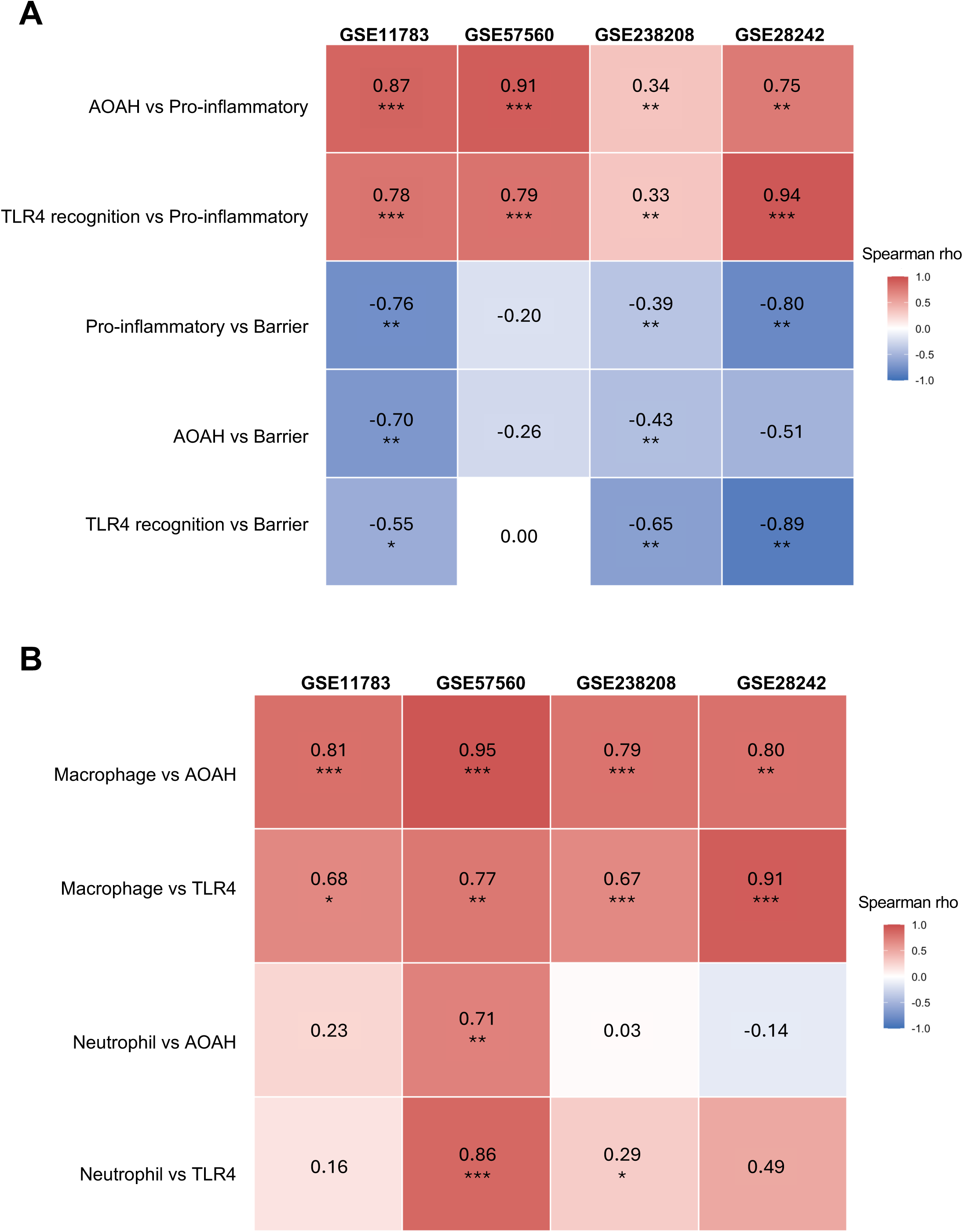
AOAH and TLR4 signaling correlate with inflammatory and immune-cell transcriptional states across IC/BPS cohorts. (A) Within-cohort associations among AOAH expression, TLR4 receptor/co-receptor recognition, pro-inflammatory signaling, and urothelial barrier integrity across GSE11783 (n=16), GSE57560 (n=16), GSE238208 (n=63), and GSE28242 (n=13). Heatmap values represent Spearman rank-correlation coefficients (ρ) calculated independently within each cohort. *AOAH* and TLR4 recognition were positively associated with pro-inflammatory signaling across cohorts, whereas inflammatory features showed inverse relationships with urothelial barrier integrity. Expanded within-cohort correlation matrices incorporating TLR4 proximal signaling, regulatory signaling, and calprotectin are shown in Supplementary Figure 14; additional cross-cohort relationships involving calprotectin are shown in Supplementary Figure 15. Individual sample-level scatterplots for the principal pathway relationships are shown in Supplementary Figure 16. (B) Associations of monocyte/macrophage- and neutrophil-associated transcriptional scores with AOAH expression and the TLR4 recognition module across the same four cohorts. Monocyte/macrophage-associated expression showed consistently strong positive correlations with both AOAH and TLR4 recognition, whereas neutrophil-associated relationships were less consistent across cohorts. Expanded immune-cell analyses, including pan-leukocyte, B-cell, T-cell, monocyte/macrophage, and neutrophil associations with AOAH and TLR4 recognition, are shown in Supplementary Figure 17. Numbers within heatmap cells indicate Spearman ρ; color indicates the direction and magnitude of the association. *adjusted p < 0.05, **adjusted p < 0.01, ***adjusted p < 0.001.

To investigate whether these molecular associations were related to immune-cell enrichment, we next correlated *AOAH* expression and TLR4 recognition scores with cell type-associated transcriptional signatures. Monocyte/macrophage signatures showed strong positive correlations with both *AOAH* and TLR4 recognition across all four cohorts (Figure 6B). In contrast, neutrophil-associated correlations were less consistent, with the strongest associations observed in GSE57560 (Figure 6B). Importantly, these relationships were not restricted to myeloid signatures. Pan-leukocyte scores correlated positively with *AOAH* and TLR4 recognition across all four cohorts, while T-cell and B-cell signatures also showed positive associations, although their strength and statistical significance varied across datasets (Supplementary Figure 17). Together, these findings indicate that elevated *AOAH* and TLR4-associated expression occur within a broader immune-enriched inflammatory transcriptional state, with particularly consistent associations involving monocyte/macrophage signatures. These correlations do not establish whether increased expression reflects immune-cell abundance, altered cellular transcriptional states, or both.

### Clinically severe IC/BPS phenotypes converge on a shared molecular state

Finally, we examined whether clinically distinct severe IC/BPS phenotypes exhibited convergent molecular remodeling. We compared ulcerative IC versus healthy controls in GSE11783 with low-capacity IC versus healthy controls in GSE57560 across nine immune, inflammatory, *AOAH*, TLR4 signaling, and urothelial features (Figure 7A). Despite independent cohort design and different clinical definitions, both severe phenotypes showed increased leukocyte and monocyte/macrophage-associated expression, increased TLR4 recognition and pro-inflammatory signaling, elevated *AOAH*, and reduced urothelial barrier-associated expression. Although the magnitude of these changes differed for some features, including neutrophil signatures and TLR4 proximal signaling, the overall effect-size profiles were strongly concordant between ulcerative and low-capacity IC (Figure 7B; Spearman ρ=0.87, p=0.0021). An expanded comparison incorporating calprotectin showed a similar pattern of shared inflammatory and urothelial transcriptional changes (Supplementary Figure 18). Together, these findings identify convergent inflammatory and urothelial transcriptional signatures across the two independently characterized severe IC/BPS phenotypes.

**Figure 7.**
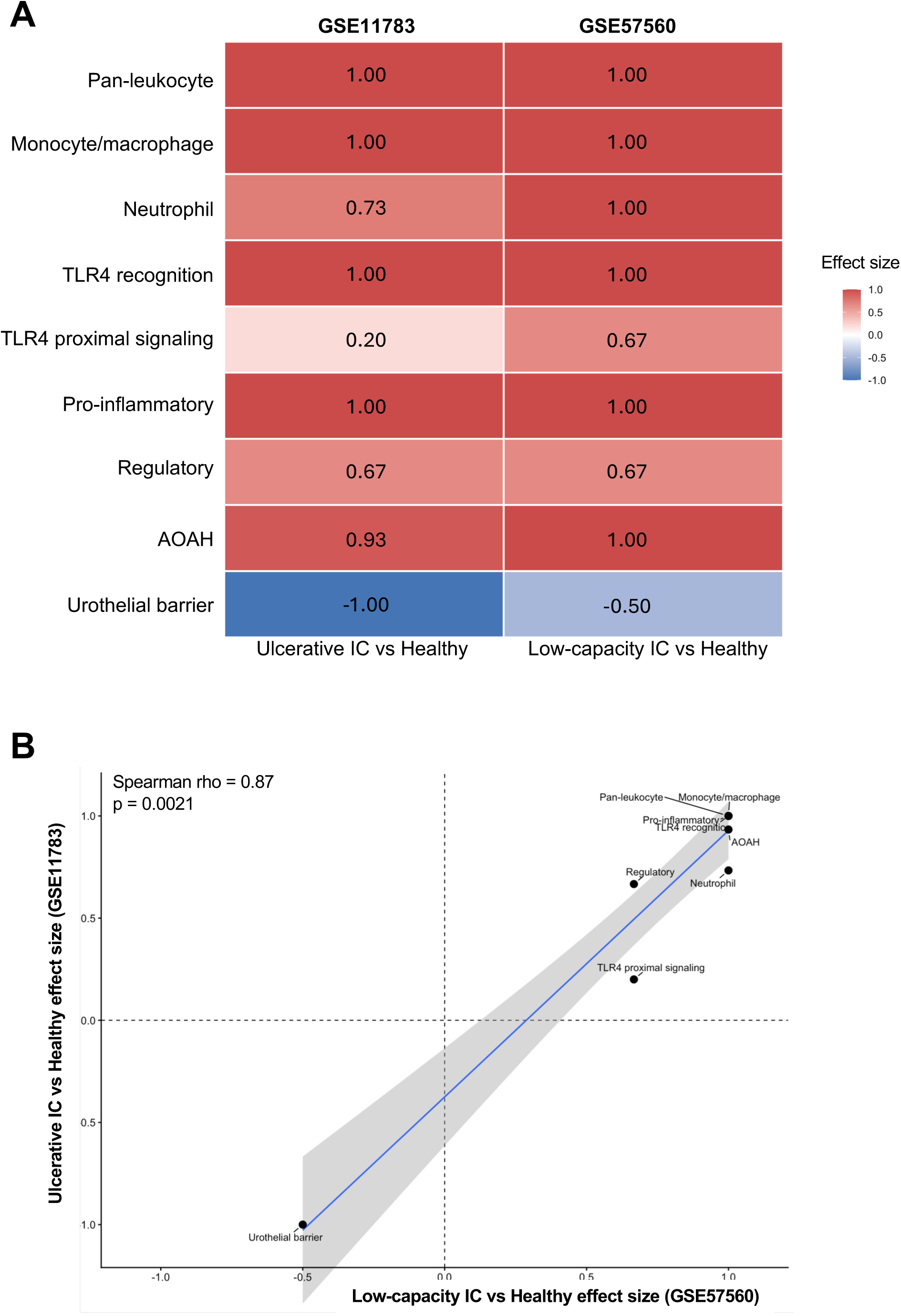
Severe IC/BPS phenotypes converge on a shared inflammatory and urothelial transcriptional state. (A) Comparison of molecular and immune-cell features associated with severe IC/BPS phenotypes in two independent bladder-tissue cohorts: ulcerative IC versus healthy controls in GSE11783 and low-capacity IC versus healthy controls in GSE57560. Heatmap values represent rank-biserial effect sizes derived from pairwise Wilcoxon rank-sum comparisons, oriented such that positive values indicate higher feature scores in the severe IC/BPS phenotype and negative values indicate lower scores relative to healthy controls. Both severe phenotypes were characterized by increased leukocyte and monocyte/macrophage signatures, TLR4 recognition and pro-inflammatory signaling, and *AOAH* expression, accompanied by reduced urothelial barrier-associated expression. The corresponding analysis including calprotectin is shown in Supplementary Figure 18. (B) Concordance of severe-phenotype effect sizes between GSE11783 and GSE57560 across the nine molecular and cellular features shown in (A). Each point represents one feature; the x-axis indicates the effect size for low-capacity IC versus healthy controls in GSE57560, and the y-axis indicates the effect size for ulcerative IC versus healthy controls in GSE11783. Effect-size profiles were strongly correlated between the two independent severe IC/BPS phenotypes (Spearman ρ = 0.87, p = 0.0021). The solid line represents a linear fit with 95% confidence interval; dashed lines indicate zero effect.

## <u>D</u>ISCUSSION

In this multi-cohort reanalysis, we identified reproducible molecular programs across four independently generated IC/BPS transcriptomic datasets despite substantial differences in clinical classification, specimen type, geography, and transcriptomic platform. Four major findings emerged. First, bladder-tissue IC/BPS was consistently associated with reduced urothelial differentiation and barrier-associated expression (Figure 3; Supplementary Figures 5-8). Second, immune remodeling was heterogeneous across disease phenotypes but repeatedly included a myeloid-associated component accompanied by increased TLR4 recognition and pro-inflammatory transcription (Figures 2 and 4). Third, *AOAH* was elevated across all three primary bladder-tissue cohorts and closely tracked monocyte/macrophage markers and inflammatory expression (Figures 5 and 6). Finally, independently defined severe IC/BPS phenotypes, specifically ulcerative and low-capacity IC, exhibited highly concordant molecular signatures (Figure 7), suggesting that clinically distinct forms of severe IC/BPS can converge on a shared inflammatory and urothelial molecular state.

A central feature of this shared molecular state is altered urothelial differentiation and barrier-associated transcription. The urothelium is not simply a passive barrier preventing the absorption of potentially harmful substances in urine; it provides a differentiated interface between urine and underlying sensory, stromal, vascular, and immune compartments [12, 38–41]. Across all three bladder-tissue datasets, IC/BPS was associated with reduced expression of genes related to urothelial differentiation and epithelial junctional integrity (Figure 3; Supplementary Figures 5-7). The recurrence of this molecular signature across non-ulcer, ulcerative, normal- and low-capacity, and background non-lesion and Hunner lesion tissue suggests that impaired urothelial differentiation is not restricted to a single clinical subtype, but that urothelial barrier dysfunction is common across all IC phenotypes [13–15, 42]. At the same time, the magnitude of barrier loss differed across groups, supporting the view that IC/BPS represents a biological spectrum rather than a uniform disease phenotype. The absence of the comparable composite signal in urine sediment also emphasizes that specimen type matters. Exfoliated urinary cells may capture inflammatory or cellular processes that are complementary to, but not equivalent to, intact bladder tissue.

Previous studies have implicated diverse immune populations in IC/BPS, including innate immune myeloid lineage cells, adaptive B and T-cells, and mast cells [16–21]. In our analysis, the immune landscape was similarly heterogeneous, but monocyte/macrophage-associated expression emerged as one of the most reproducible features (Figure 2). T- and B-cell signatures were also elevated in selected phenotypes, consistent with evidence for adaptive immune involvement in IC/BPS, particularly in more inflammatory sub-phenotypes. Notably, monocyte/macrophage scores showed especially strong associations with both *AOAH* and TLR4 recognition across cohorts (Figure 6B). However, pan-leukocyte, T-cell, and B-cell signatures also showed positive associations with these molecular features (Supplementary Figure 17), indicating that the relationships were not restricted to myeloid-associated transcription. Thus, while immune-cell composition may vary across clinical phenotypes, broader immune enrichment, including a consistent monocyte/macrophage-associated component, is a shared feature of these disparate IC/BPS cohorts. These findings also parallel observations in chronic prostatitis, where monocyte/macrophage-associated inflammation has been implicated in disease pathophysiology [39,40]. Clinical studies have identified monocyte/macrophage populations associated with inflammatory markers, while experimental prostatitis models demonstrate macrophage recruitment near sensory afferents and suggest a functional contribution to chronic pelvic pain [43–45].

TLR4 signaling provides a plausible mechanistic link between this myeloid inflammatory state and prior microbial exposure. As recently discussed, microbial exposures and host-microbiota interactions may contribute to IC/BPS pathophysiology, providing a framework for investigating innate immune mechanisms in persistent bladder pain [22]. In fact, studies from the MAPP Research Network have previously implicated TLR4 signaling with IC/BPS pain in specific phenotypes [23–25]. The recognition module evaluated here includes genes involved in TLR4-mediated LPS recognition, including *TLR4*, *CD14, LY96,* and *LBP,* and was increased across the primary bladder-tissue cohorts together with downstream inflammatory transcription (Figure 4). These bulk transcriptomic data cannot establish receptor activation or identify the initiating ligand. Nevertheless, the reproducibility of the TLR4-associated signal across independent populations supports prior work implicating innate immune sensing in IC/BPS and is compatible with models in which infection or microbial products initiate or perpetuate inflammatory responses in a susceptible host. This interpretation is particularly relevant to post-UTI pelvic pain models, in which transient infection with O-antigen-deficient *E. coli* isolates induces persistent pain and urinary phenotypes after apparent bacterial clearance [46–50]. To explore whether similar molecular alterations occur following urinary tract infection, we compared the human IC/BPS transcriptional profiles with bladder transcriptomic data from the post-UTI pelvic pain (PUPP) mouse model (Supplementary Figure 19). This comparison revealed shared directional changes in several disease-associated features, most notably reduced expression of urothelial differentiation and barrier-related genes. Although inflammatory transcriptional changes were less consistently conserved across species, these findings suggest that prior urinary infection can produce aspects of the urothelial molecular phenotype observed in human IC/BPS and is consistent with the reported UTI history among IC/BPS patients [51].

AOAH may be considered in relation to TLR4 signaling rather than as a canonical signaling component upstream or downstream of TLR4. AOAH, a hydrolase, cleaves secondary acyl chains from hexa-acyl LPS into a tetra-acyl form, thereby changing a stimulatory molecule into an inhibitor, hence limiting prolonged LPS bioactivity and decreasing TLR4-dependent inflammation and tissue death [52]. At the same time, AOAH is strongly associated with myeloid cells. The marked correlation between *AOAH* and monocyte/macrophage signatures across all four cohorts therefore likely reflects, at least in part, the cellular context in which AOAH is expressed (Figure 6B). However, growing evidence from human genetic studies suggests that AOAH may also function as a biological modifier rather than merely a marker of myeloid abundance [29–33]. Together with prior genetic associations and experimental findings in AOAH-deficient models [34–37], our analyses here support a TLR4-AOAH axis in which inflammatory LPS sensing and its regulation intersect with myeloid remodeling in IC/BPS. The present transcriptomic study establishes association within this axis but cannot distinguish between increased AOAH as compensatory induction, altered cellular composition, intrinsic myeloid response, or a combination of these mechanisms.

The inverse relationships between inflammatory features and urothelial barrier expression further suggest that epithelial dysfunction and inflammation are coordinated aspects of the IC/BPS molecular state. Across multiple cohorts, higher TLR4 pro-inflammatory transcription and *AOAH* expression accompanied lower urothelial barrier scores (Figure 6A; Supplementary Figure 16). The directionality of this relationship cannot be determined from cross-sectional transcriptomic data. It is plausible that impaired urothelial integrity could increase exposure of the underlying bladder wall to urinary or microbial products, thereby promoting immune recruitment and activation. Conversely, inflammatory mediators could disrupt urothelial differentiation and junctional integrity. These mechanisms are not mutually exclusive and may form a self-reinforcing vicious cycle of epithelial dysfunction and inflammation [12, 13]. Importantly, the attenuation of the relationship between *AOAH* and urothelial barrier expression following adjustment for immune-cell signatures suggests that *AOAH* may primarily reflect the broader inflammatory tissue environment rather than directly mediate epithelial dysfunction.

The findings here further support IC/BPS as a two-hit or multi-hit disease. For example, microbial exposure such as UTI could provide an initiating inflammatory stimulus, while inherent host factors, including genetic predisposition or variants that affect innate immune regulation, could influence whether that response resolves or becomes persistent. Experimental studies have demonstrated that transient bacterial infection can induce chronic pelvic pain after bacterial clearance and that susceptibility depends on both bacterial characteristics and host genetic background [46, 53]. Furthermore, studies in mice have identified AOAH as a genetic modulator of pelvic pain severity [37]. Together, these observations suggest that different combinations of epithelial vulnerability, immune composition, prior inflammatory exposures, and host genetic background may contribute to clinical heterogeneity while converging on shared molecular programs.

The strong concordance between ulcerative and low-capacity disease is particularly notable in this context. These clinical sub-phenotypes were defined using different diagnostic and phenotypic clinical criteria, yet their molecular effect-size profiles across nine selected features were highly correlated (Figure 7). Both phenotypes exhibited increased myeloid-associated and inflammatory transcription, including TLR4 recognition and *AOAH* expression, accompanied by reduced urothelial barrier-associated expression. This convergence suggests that molecular state may provide information not captured by existing clinical labels alone. Conversely, the differences observed between less severe and severe subgroups reinforce the importance of patient phenotyping in both mechanistic studies and clinical trials. Molecular characterization may ultimately complement clinical classification by identifying patients with prominent epithelial, myeloid-inflammatory, or mixed disease features.

This retrospective analysis relied on publicly available datasets with heterogeneous clinical classifications, specimen types, and transcriptomic platforms, with relatively small sample sizes in some cohorts, thus posing limitations (Table 1). Bulk transcriptomic data cannot distinguish changes in immune-cell abundance from cell-intrinsic transcriptional responses, highlighting the need for future single-cell and spatially resolved analyses. Additionally, the BCG-treated comparator group in GSE238208 is biologically distinct from healthy controls, and the cross-sectional design precludes causal inference. Future studies integrating larger, clinically well-characterized cohorts with single-cell and spatial transcriptomics, longitudinal sampling, and functional validation will be important for defining the cellular and mechanistic contributions of the TLR4-AOAH axis to IC/BPS.

Despite the limitations of heterogeneous clinical and molecular methodologies, the convergence of molecular findings across independent cohorts is a major strength of this study. By analyzing each cohort independently and identifying recurrent features across heterogeneous clinical populations, we demonstrate a shared IC/BPS-associated molecular architecture characterized by reduced urothelial differentiation, myeloid-associated immune remodeling, increased TLR4-associated inflammatory transcription, and elevated *AOAH* expression. The striking concordance between independently defined severe IC/BPS phenotypes further suggests that distinct clinical presentations and/or etiologies may converge on common molecular states. Together, these findings provide a human transcriptomic framework for investigating the TLR4-AOAH axis as a potential link between innate immune regulation, urothelial dysfunction, and persistent bladder pain. Ultimately, integrating molecular signatures with clinical phenotyping and mechanistic studies may help identify biologically distinct patient populations and inform more targeted therapeutic strategies for IC/BPS.

## MATERIALS AND METHODS

### Dataset identification and study design

Publicly available transcriptomic datasets from patients with interstitial cystitis/bladder pain syndrome (IC/BPS) were obtained from Gene Expression Omnibus (GEO). Four independent cohorts were included: GSE11783, GSE57560, GSE238208, and GSE28242 (Table 1; Fig. 1). GSE11783 comprised bladder-biopsy microarray samples from healthy controls (n=6), non-ulcer IC (n=5), and ulcerative IC (n=5). GSE57560 comprised bladder-biopsy microarray samples from healthy controls (n=3), normal-capacity IC (n=9), and low-capacity IC (n=4). GSE238208 comprised bladder biopsy samples from patients with Hunner-type IC, including matched lesion and non-lesion specimens from 25 patients, together with 13 Bacillus Calmette-Guérin (BCG)-associated cystitis comparator samples. GSE28242 comprised urine-sediment microarray samples from healthy controls (n=5), non-lesion IC (n=5), and lesion IC (n=3). The three bladder-tissue cohorts were used to investigate disease-associated transcriptional alterations, while the urine sediment cohort provided an independent specimen-based comparison. All cohorts were analyzed separately to account for differences in study design, sample source, and expression platform.

### Transcriptomic preprocessing

Expression data and accompanying sample metadata were obtained from GEO. Microarray probes were mapped to gene symbols to generate gene-level expression matrices. Representative probe profiles were retained for genes represented by multiple probes. For GSE238208, the available processed RNA-sequencing expression matrix was used. Analyses were performed independently within each cohort without pooling expression values across platforms.

### Definition of cell-type and molecular transcriptional features

Predefined marker sets were used to estimate tissue-compartment, immune-cell, epithelial, and inflammatory transcriptional states (Table 2). Cell-associated features included general urothelium (*EPCAM, KRT7, KRT8, KRT18, KRT19*); basal/regenerative urothelium (*KRT5, KRT14, KRT17, TP63*); smooth muscle (*ACTA2, TAGLN, MYH11, CNN1, DES*); fibroblast/stroma (*COL1A1, COL1A2, COL3A1, DCN, LUM, COL6A1*); endothelium (*PECAM1, VWF, KDR, EMCN, ENG*); pan-leukocytes (*PTPRC, CORO1A, LST1, TYROBP, FCER1G*); monocytes/macrophages (*LST1, TYROBP, FCER1G, AIF1, CD68, CTSS*); neutrophils (*FCGR3B, CSF3R, CEACAM8, MPO, ELANE, CTSG*); T cells (*CD3D, CD3E, CD3G, TRAC, TRBC1*); B cells (*CD79A, CD79B, MS4A1, CD37, CD74*); and mast cells (*TPSAB1, TPSB2, CPA3, KIT, MS4A2, HDC*). The urothelial barrier module comprised differentiation genes (*UPK1A, UPK1B, UPK2, UPK3A, KRT20*) and tight junction/barrier genes (*TJP1, OCLN, CLDN4, CLDN8*). The TLR4 recognition module comprised *TLR4, CD14, LY96,* and *LBP*; the proximal TLR4 signaling module comprised *TIRAP, MYD88,* and *TICAM1*; the pro-inflammatory module comprised *IL1B, IL6, TNF, CCL2, CXCL8, PTGS2,* and *MMP9*; and the regulatory module comprised *IL4, IL4R, IL10, IL10RA, STAT6, SOCS3,* and *TGFB1*. *AOAH* was analyzed individually. Calprotectin-associated transcription was represented by *S100A8* and *S100A9*.

### Calculation of transcriptional module scores

Transcriptional module scores were calculated independently within each cohort. For each gene, expression values were standardized across samples by subtracting the cohort-specific mean and dividing by the standard deviation. For each sample, a module score was calculated as the arithmetic mean of the standardized expression values for the genes included in that module. Genes absent from a dataset or lacking sufficient expression variability were excluded from the corresponding module calculation. This approach generated relative transcriptional feature scores within each cohort while reducing differences in expression magnitude among genes.

### Group comparisons and effect-size estimation

Differences in transcriptional feature scores among clinical groups were assessed independently within each cohort using Kruskal–Wallis tests, followed by Dunn’s post hoc comparisons with Benjamini–Hochberg adjustment for multiple testing. Pairwise effect sizes were estimated using rank-biserial correlations derived from Wilcoxon rank-sum statistics. Comparisons were performed relative to healthy controls in GSE11783, GSE57560, and GSE28242, and relative to BCG-associated cystitis comparator samples in GSE238208. Because GSE238208 contained matched Hunner lesion and non-lesion biopsies from the same patients, an additional paired analysis was performed using Wilcoxon signed-rank tests, with Benjamini–Hochberg adjustment across the 11 cell-type transcriptional features.

### Correlation analyses

Associations among transcriptional features were evaluated independently within each cohort using Spearman’s rank correlation coefficients (ρ). Correlations were examined among AOAH expression, TLR4-associated signaling modules, inflammatory signaling, urothelial barrier scores, and inferred immune-cell transcriptional features. P-values were adjusted for multiple comparisons using the Benjamini–Hochberg method, and adjusted P-values were used to assign significance annotations in correlation heatmaps.

### Cross-cohort comparison of severe IC/BPS phenotypes

To investigate shared transcriptional alterations across clinically severe IC/BPS phenotypes, effect sizes were compared between ulcerative IC versus healthy controls in GSE11783 and low-capacity IC versus healthy controls in GSE57560. Nine transcriptional features were included: pan-leukocyte, monocyte/macrophage, neutrophil, TLR4 recognition, TLR4 proximal signaling, pro-inflammatory signaling, regulatory signaling, *AOAH*, and urothelial barrier function. Concordance in effect sizes between the two comparisons was assessed using Spearman’s rank correlation (ρ). An expanded analysis incorporating calprotectin as the tenth feature was performed separately (Supplementary Figure 18).

### Statistical analysis and visualization

All analyses were performed in R version 4.4.1 using packages including *dplyr* (1.2.0), *tidyr* (1.3.2), *rstatix* (0.7.3), *ggplot2* (4.0.2), *ggpubr* (0.6.3), and *patchwork* (1.3.2). Boxplots display medians and interquartile ranges, with whiskers extending to 1.5 times the interquartile range. Statistical significance was defined as an adjusted P-value below 0.05 unless otherwise indicated.

### Cross-species comparison with the post-UTI pelvic pain model

To investigate whether transcriptional alterations observed in human IC/BPS were also present following urinary tract infection, we performed an exploratory cross-species comparison using bladder gene-expression data from the mouse post-UTI pelvic pain (PUPP) model (Supplementary Figure 19). Expression profiles from vehicle-treated mice (n = 4) and mice exposed to uropathogenic *Escherichia coli* NU23 (n = 4) were analyzed. Genes with duplicate symbols were collapsed by averaging their expression values across samples. Expression values were log2-transformed after addition of a pseudocount of 1 and standardized for each gene across samples using z-scores.

Six transcriptional features were examined: *AOAH*, TLR4 recognition, TLR4 proximal signaling, pro-inflammatory signaling, regulatory signaling, and urothelial barrier-associated expression. Mouse gene sets were defined using the corresponding murine gene symbols. For each sample, module scores were calculated as the mean standardized expression of available genes within each feature. Differences between NU23-exposed and vehicle-treated mice were evaluated using Wilcoxon rank-sum tests, with Benjamini–Hochberg correction across the six features.

For comparison with human IC/BPS, corresponding transcriptional features were examined in ulcerative IC versus healthy controls (GSE11783), low-capacity IC versus healthy controls (GSE57560), Hunner lesion tissue versus BCG-associated cystitis comparators (GSE238208), and lesion IC versus healthy controls (GSE28242). Cross-species concordance was visualized using a heatmap displaying the fraction of genes within each feature showing increased expression in the disease-associated group relative to its comparator. Significance annotations reflected adjusted P-values from the corresponding within-cohort module comparisons. Absolute expression values were not compared directly between species.

---

**Supplementary Figure 1. Cell-type and tissue-compartment transcriptional signatures across IC/BPS cohorts in GSE11783.** Boxplots show urothelial, endothelial, stromal/smooth-muscle, and immune-cell signature scores across clinical sub-phenotypes in GSE11783, illustrating cohort-specific patterns of tissue remodeling and immune-cell enrichment.

**Supplementary Figure 2. Cell-type and tissue-compartment transcriptional signatures across IC/BPS cohort in GSE57560.** Boxplots show urothelial, endothelial, stromal/smooth-muscle, and immune-cell signature scores across clinical sub-phenotypes in GSE57560, illustrating cohort-specific patterns of tissue remodeling and immune-cell enrichment.

**Supplementary Figure 3. Cell-type and tissue-compartment transcriptional signatures across IC/BPS cohort in GSE238208.** Boxplots show urothelial, endothelial, stromal/smooth-muscle, and immune-cell signature scores across clinical sub-phenotypes in GSE238208, illustrating cohort-specific patterns of tissue remodeling and immune-cell enrichment.

**Supplementary Figure 4. Cell-type and tissue-compartment transcriptional signatures across IC/BPS cohort in GSE28242.** Boxplots show urothelial, endothelial, stromal/smooth-muscle, and immune-cell signature scores across clinical sub-phenotypes in GSE28242, illustrating cohort-specific patterns of tissue remodeling and immune-cell enrichment.

**Supplementary Figure 5. Individual urothelial differentiation and epithelial barrier gene expression in GSE11783.** Boxplots show expression of UPK1A, UPK1B, UPK2, UPK3A, KRT20, TJP1, OCLN, CLDN4, and CLDN8 across clinical sub-phenotypes in GSE11783, providing gene-level support for the urothelial barrier module shown in Figure 3.

**Supplementary Figure 6. Individual urothelial differentiation and epithelial barrier gene expression in GSE57560.** Boxplots show expression of UPK1A, UPK1B, UPK2, UPK3A, KRT20, TJP1, OCLN, CLDN4, and CLDN8 across clinical sub-phenotypes in GSE57560, providing gene-level support for the urothelial barrier module shown in Figure 3.

**Supplementary Figure 7. Individual urothelial differentiation and epithelial barrier gene expression in GSE238208.** Boxplots show expression of UPK1A, UPK1B, UPK2, UPK3A, KRT20, TJP1, OCLN, CLDN4, and CLDN8 across clinical sub-phenotypes in GSE238208, providing gene-level support for the urothelial barrier module shown in Figure 3.

**Supplementary Figure 8. Individual urothelial differentiation and epithelial barrier gene expression in GSE28242.** Boxplots show expression of UPK1A, UPK1B, UPK2, UPK3A, KRT20, TJP1, OCLN, CLDN4, and CLDN8 across clinical sub-phenotypes in GSE28242, providing gene-level support for the urothelial barrier module shown in Figure 3.

**Supplementary Figure 9. TLR4-associated inflammatory signaling in the GSE28242 cohort.** Boxplots show (A) TLR4 recognition and (B) pro-inflammatory module scores across healthy, non-lesion IC, and lesion IC samples.

**Supplementary Figure 10. TLR4 proximal signaling across IC/BPS cohorts.** Boxplots show TLR4 proximal signaling module scores across clinical sub-phenotypes in (A) GSE11783, (B) GSE57560, (C) GSE238208, and (D) GSE28242, demonstrating cohort-dependent alterations in downstream TLR4 signaling.

**Supplementary Figure 11. Regulatory signaling across IC/BPS cohorts.** Boxplots show regulatory module scores across clinical sub-phenotypes in (A) GSE11783, (B) GSE57560, (C) GSE238208, and (D) GSE28242, illustrating cohort-dependent variation in regulatory inflammatory signaling.

**Supplementary Figure 12. AOAH expression in the GSE28242 cohort.** (A) Boxplots show AOAH expression across healthy, non-lesion IC, and lesion IC samples, demonstrating phenotype-associated variation in AOAH expression.

**Supplementary Figure 13. Calprotectin-associated transcriptional signatures across IC/BPS cohorts.** Boxplots show calprotectin module scores across clinical sub-phenotypes in (A) GSE11783, (B) GSE57560, (C) GSE238208, and (D) GSE28242, demonstrating cohort- and phenotype-dependent variation in calprotectin-associated inflammatory signaling.

**Supplementary Figure 14. Correlation structure of inflammatory, AOAH–TLR4, calprotectin, and urothelial barrier signatures across IC/BPS cohorts.** Spearman correlation matrices show relationships among AOAH, TLR4 recognition and proximal signaling, pro-inflammatory and regulatory signaling, calprotectin, and urothelial barrier scores in (A) GSE11783, (B) GSE57560, (C) GSE238208, and (D) GSE28242, with correlation coefficients and statistical significance indicated within each matrix.

**Supplementary Figure 15. Calprotectin-inclusive cross-cohort pathway relationships in IC/BPS.** (A) Spearman correlations between calprotectin module scores and urothelial barrier, pro-inflammatory, and TLR4 recognition programs across the four IC/BPS cohorts. (B) Cross-cohort Spearman correlation matrix summarizing relationships among AOAH, TLR4 recognition, pro-inflammatory signaling, calprotectin, and urothelial barrier programs, extending the main-figure analysis to include calprotectin. *p < 0.05, **p < 0.01, ***p < 0.001.

**Supplementary Figure 16. Cohort-specific relationships among AOAH, inflammatory signaling, calprotectin, and urothelial barrier programs in IC/BPS.** Scatterplots show Spearman correlations across the four IC/BPS cohorts between (A) AOAH expression and urothelial barrier scores, (B) AOAH expression and pro-inflammatory module scores, (C) pro-inflammatory and urothelial barrier scores, (D) TLR4 recognition and pro-inflammatory module scores, (E) AOAH expression and calprotectin module scores, (F) calprotectin and pro-inflammatory module scores, and (G) calprotectin and urothelial barrier scores. Points represent individual samples and are colored by clinical subgroup; lines indicate linear fits with 95% confidence intervals, while inset values report Spearman ρ, p values, and sample sizes.

**Supplementary Figure 17. Immune-cell associations with AOAH expression and TLR4-associated signaling across IC/BPS cohorts. (**A) Spearman correlation heatmap showing associations of pan-leukocyte, B-cell, T-cell, monocyte/macrophage, and neutrophil signature scores with AOAH expression and TLR4 recognition module scores across GSE11783, GSE57560, GSE238208, and GSE28242. Values indicate Spearman ρ; *\**p < 0.05, **p < 0.01, ***p < 0.001.

**Supplementary Figure 18. Calprotectin-inclusive comparison of shared molecular and immune features across severe IC/BPS phenotypes.** (A) Heatmap comparing effect sizes for immune-cell signatures, TLR4 recognition and proximal signaling, pro-inflammatory and regulatory programs, AOAH, calprotectin, and the urothelial barrier in ulcerative IC versus healthy controls (GSE11783) and low-capacity IC versus healthy controls (GSE57560). Positive values indicate enrichment in the severe IC/BPS phenotype, whereas negative values indicate relative depletion; asterisks denote statistically significant differences.

**Supplementary Figure 19. Cross-species directional concordance of IC/BPS-associated transcriptional programs.** Heatmap shows the fraction of genes within each pre-specified module that are upregulated in PUPP bladder tissue and in severe human IC/BPS phenotypes relative to their respective controls. Red indicates predominantly upregulated genes, blue indicates predominantly downregulated genes, and white indicates mixed directionality. Asterisks denote BH-adjusted significance for the corresponding severe phenotype versus control comparison (*p_adj_ < 0.05, **p_adj_ < 0.01, ***p_adj_ < 0.001). Human comparisons are ulcerative IC vs healthy, low-capacity IC vs healthy, Hunner IC vs BCG, and lesion IC vs healthy. PUPP compares NU23-instilled vs vehicle (PBS-instilled) 8-week-old females; n = 4/group, and no PUPP module remained significant after multiple-testing correction

