## Supplementary Figs 1-19 for "Multi-Cohort Transcriptomic Analysis Reveals Shared Inflammatory Signaling and Urothelial Dysfunction Across IC/BPS Phenotypes"

**Supplementary Figure 19. Cross-species directional concordance of IC/BPS-associated transcriptional programs.** Heatmap shows the fraction of genes within each pre-specified module that are upregulated in PUPP bladder tissue and in severe human IC/BPS phenotypes relative to their respective controls. Red indicates predominantly upregulated genes, blue indicates predominantly downregulated genes, and white indicates mixed directionality. Asterisks denote BH-adjusted significance for the corresponding severe phenotype versus control comparison (*$p_{adj} < 0.05$, **$p_{adj} < 0.01$, ***$p_{adj} < 0.001$). Human comparisons are ulcerative IC vs healthy, low-capacity IC vs healthy, Hunner IC vs BCG, and lesion IC vs healthy. PUPP compares NU23-instilled vs vehicle (PBS-instilled) 8-week-old females; n = 4/group, and no PUPP module remained significant after multiple-testing correction.

### Supplementary Figure 1.

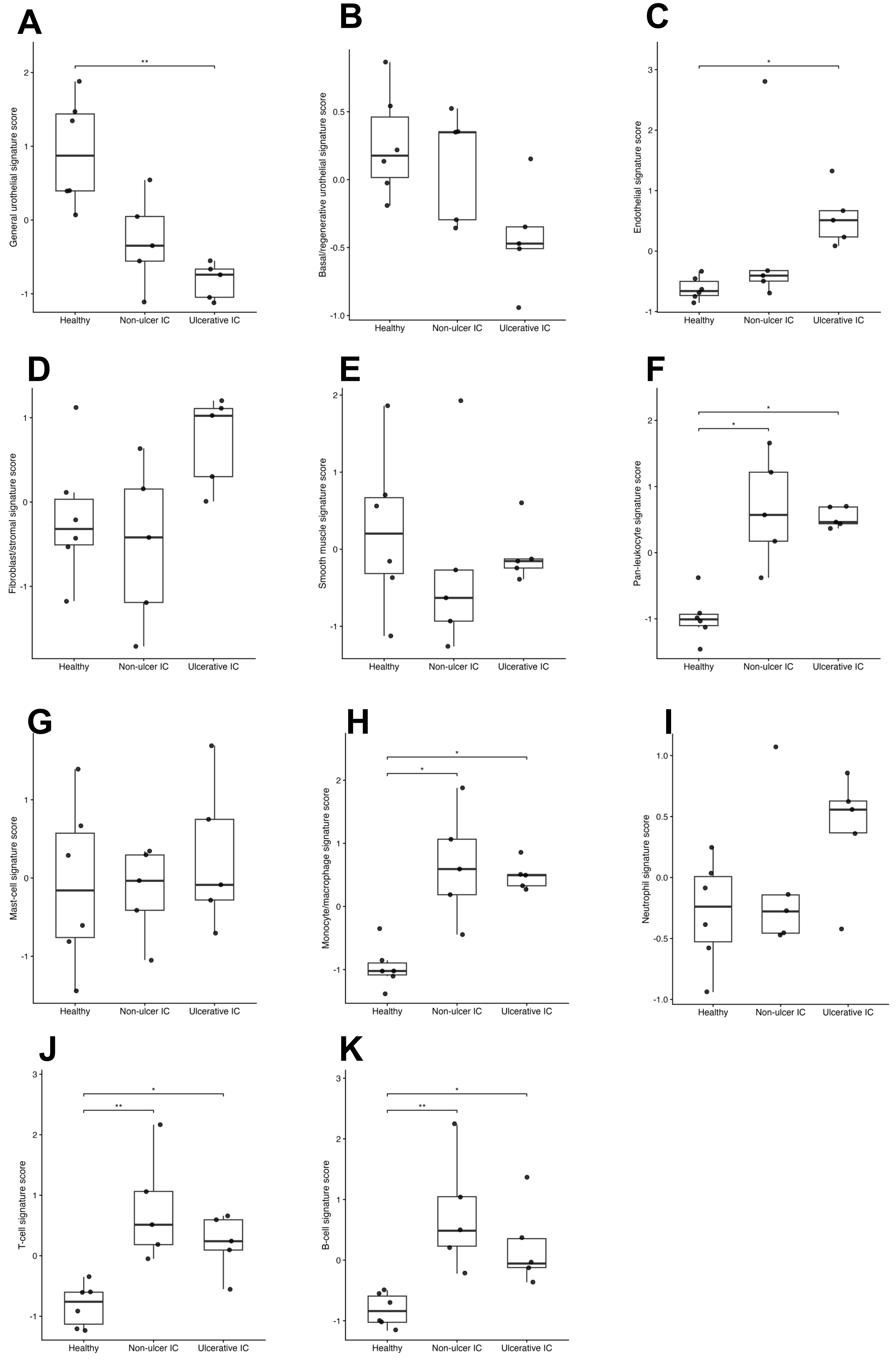

### Supplementary Figure 2.

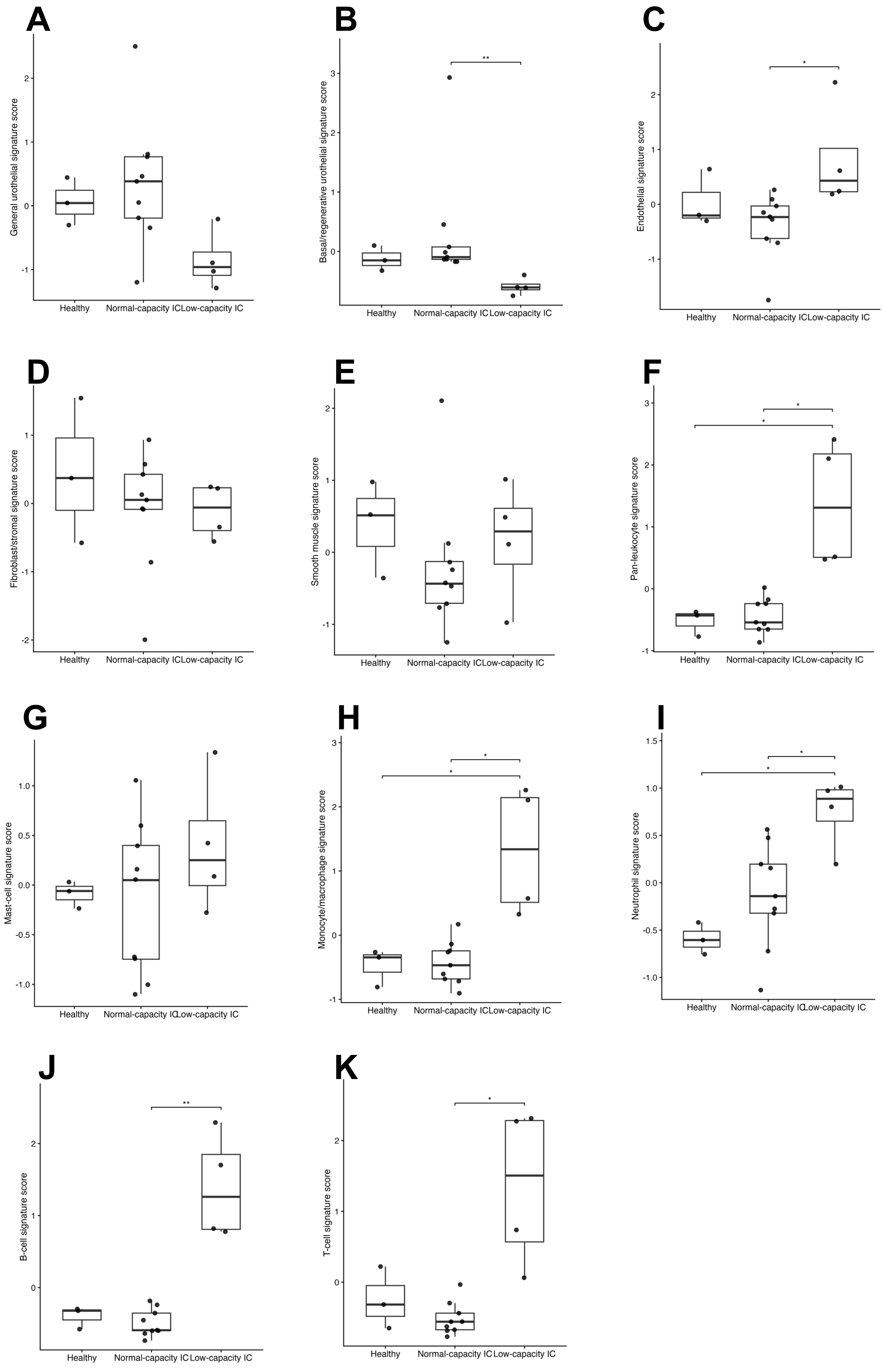

### Supplementary Figure 3.

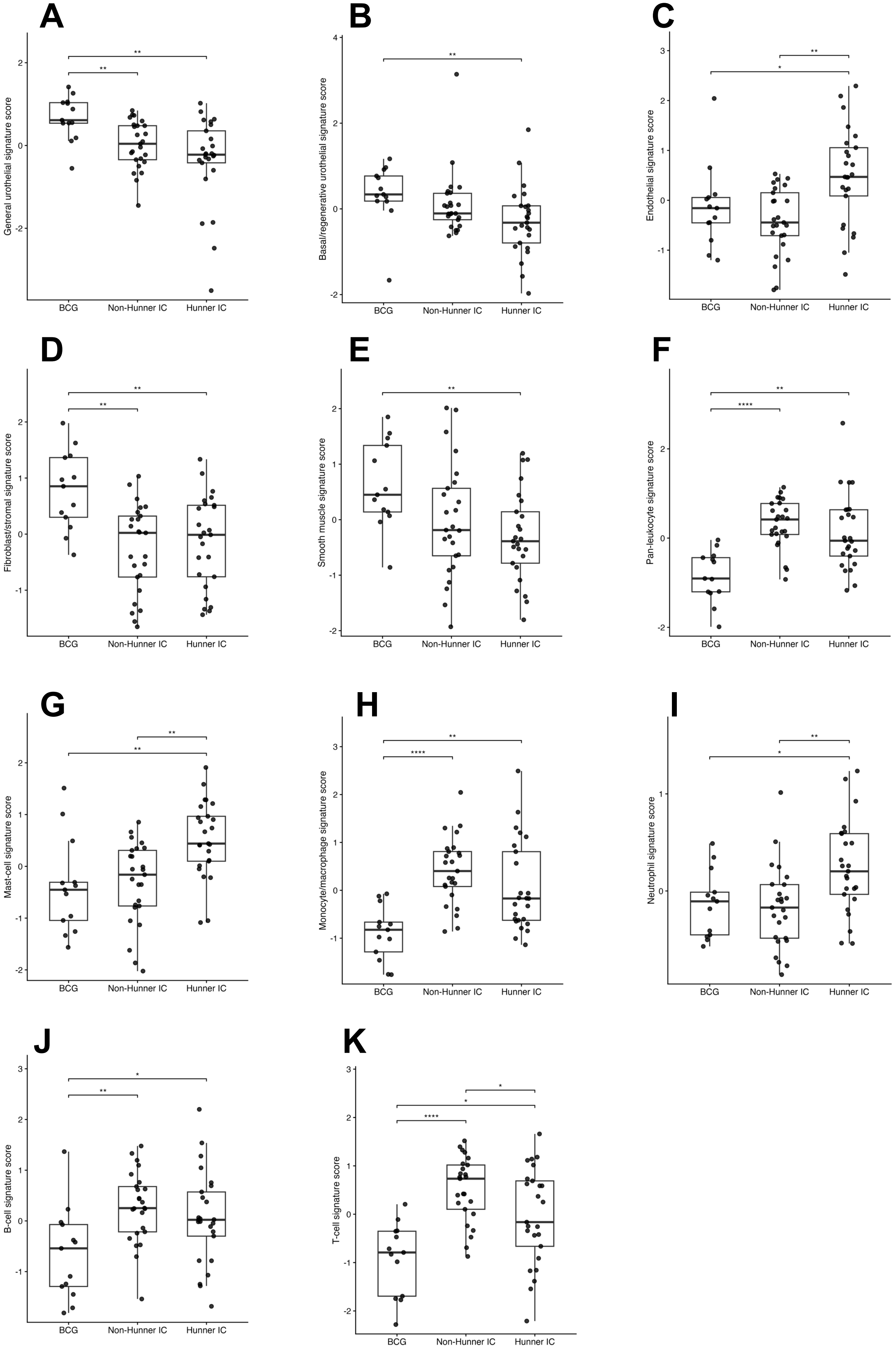

### Supplementary Figure 4.

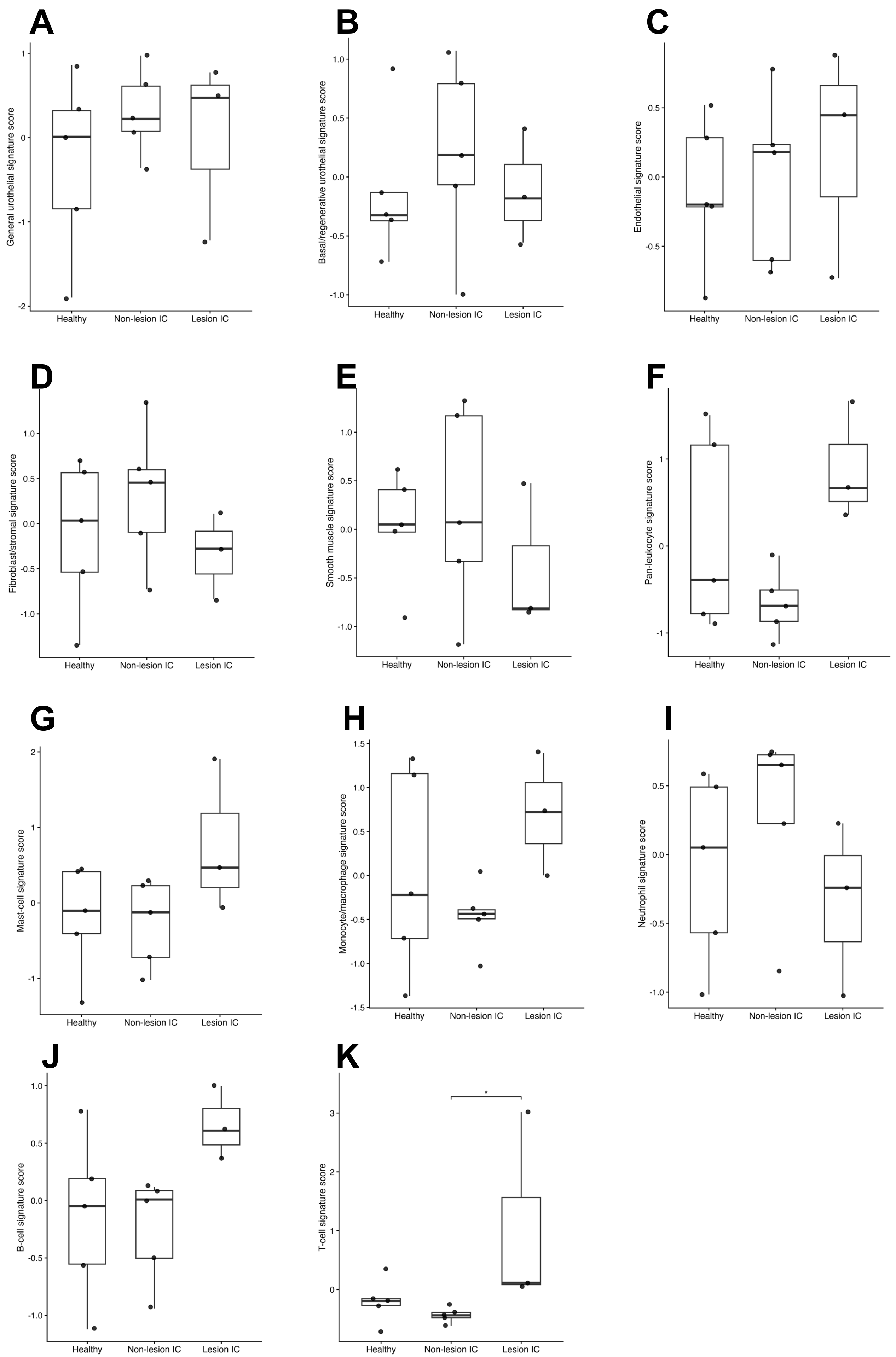

### Supplementary Figure 5.

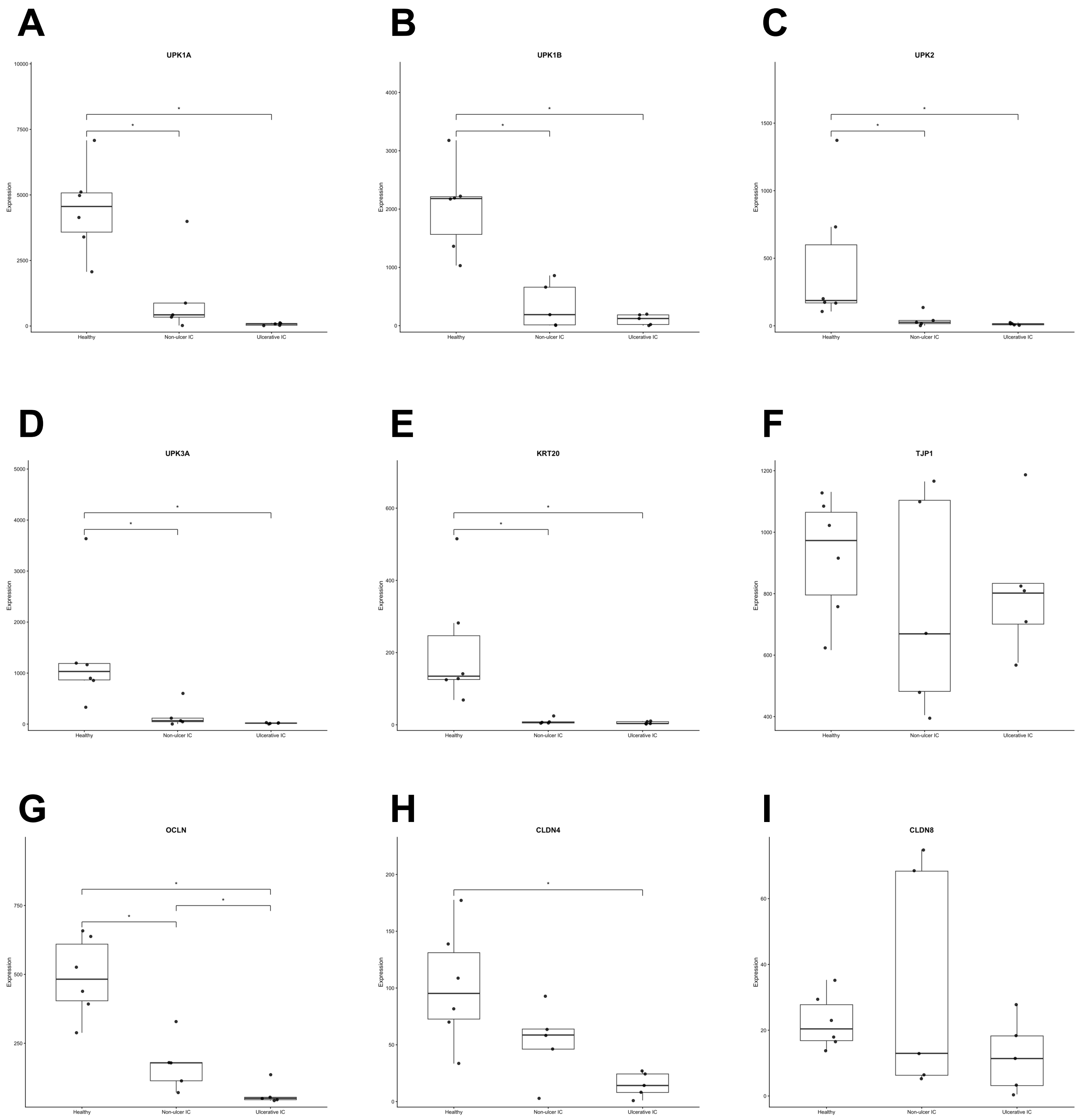

### Supplementary Figure 6.

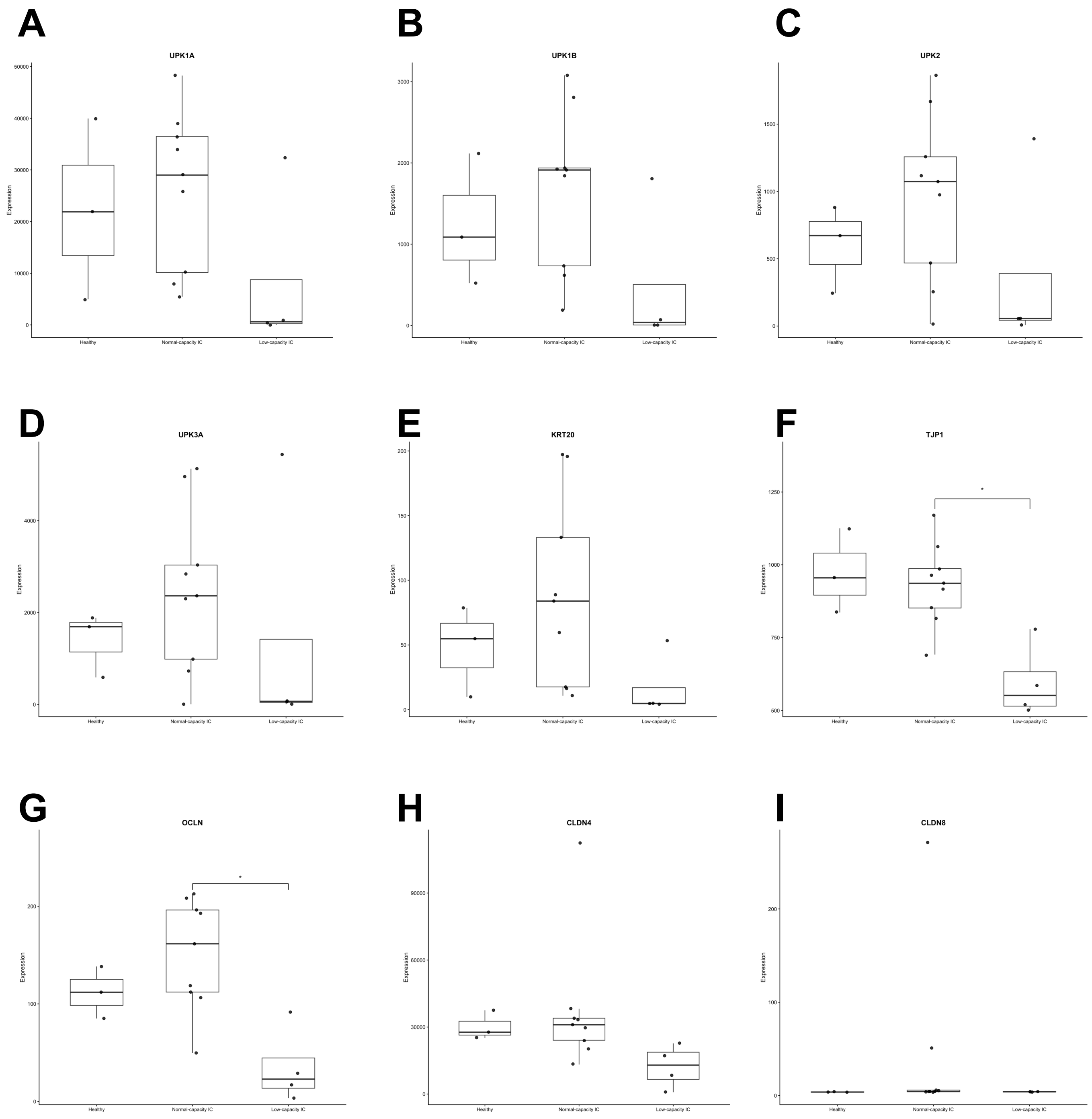

### Supplementary Figure 7.

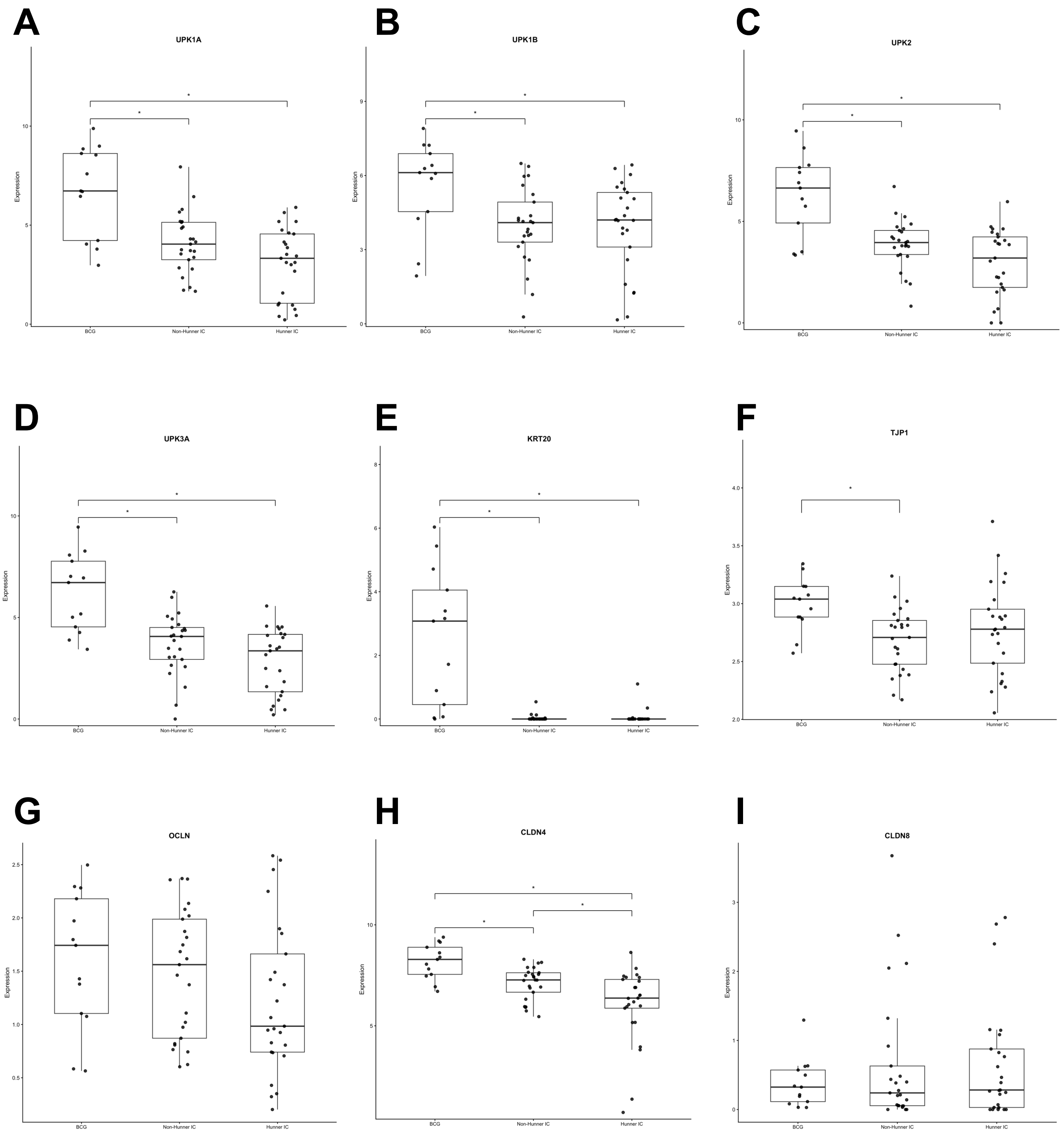

### Supplementary Figure 8.

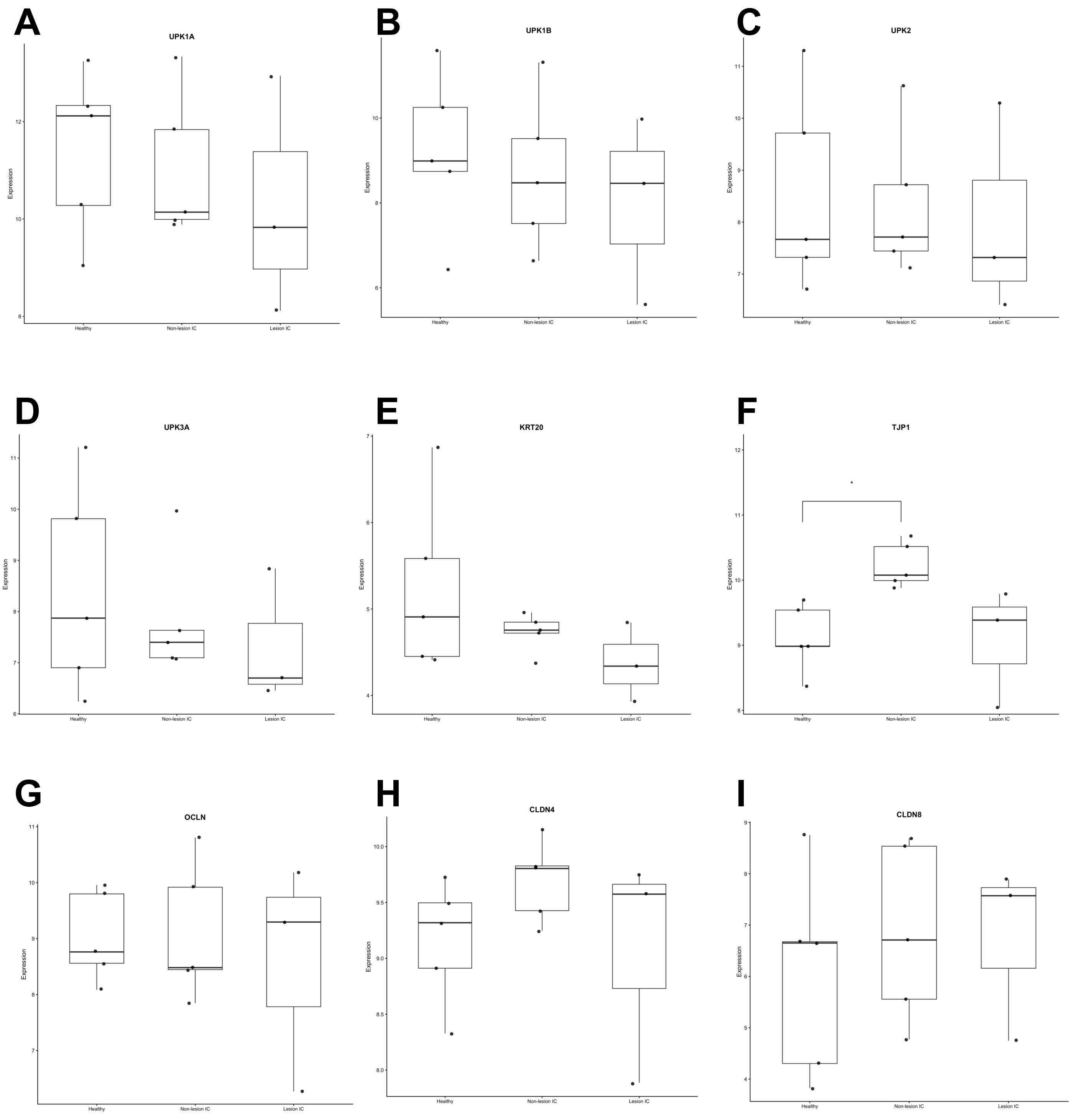

### Supplementary Figure 9.

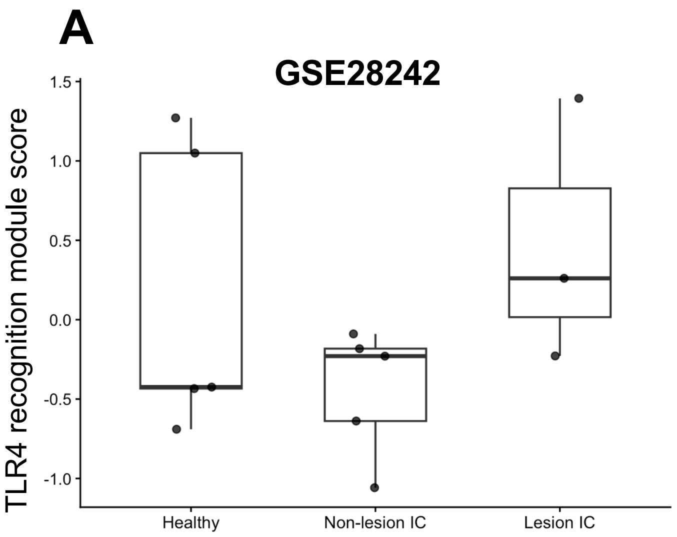


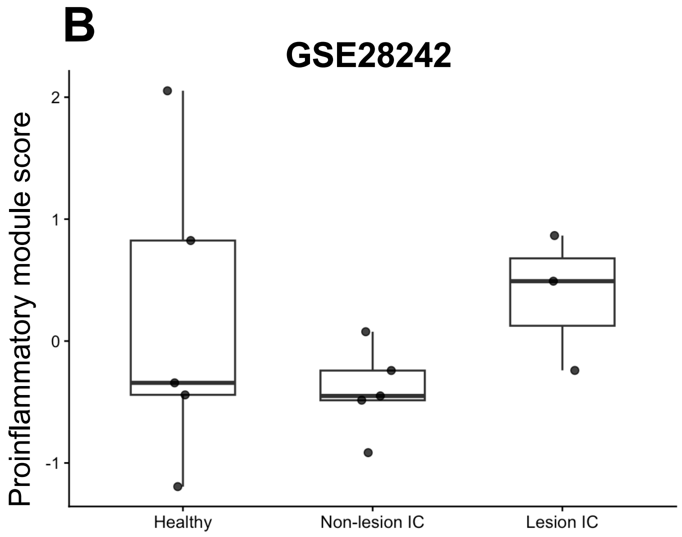

### Supplementary Figure 10.

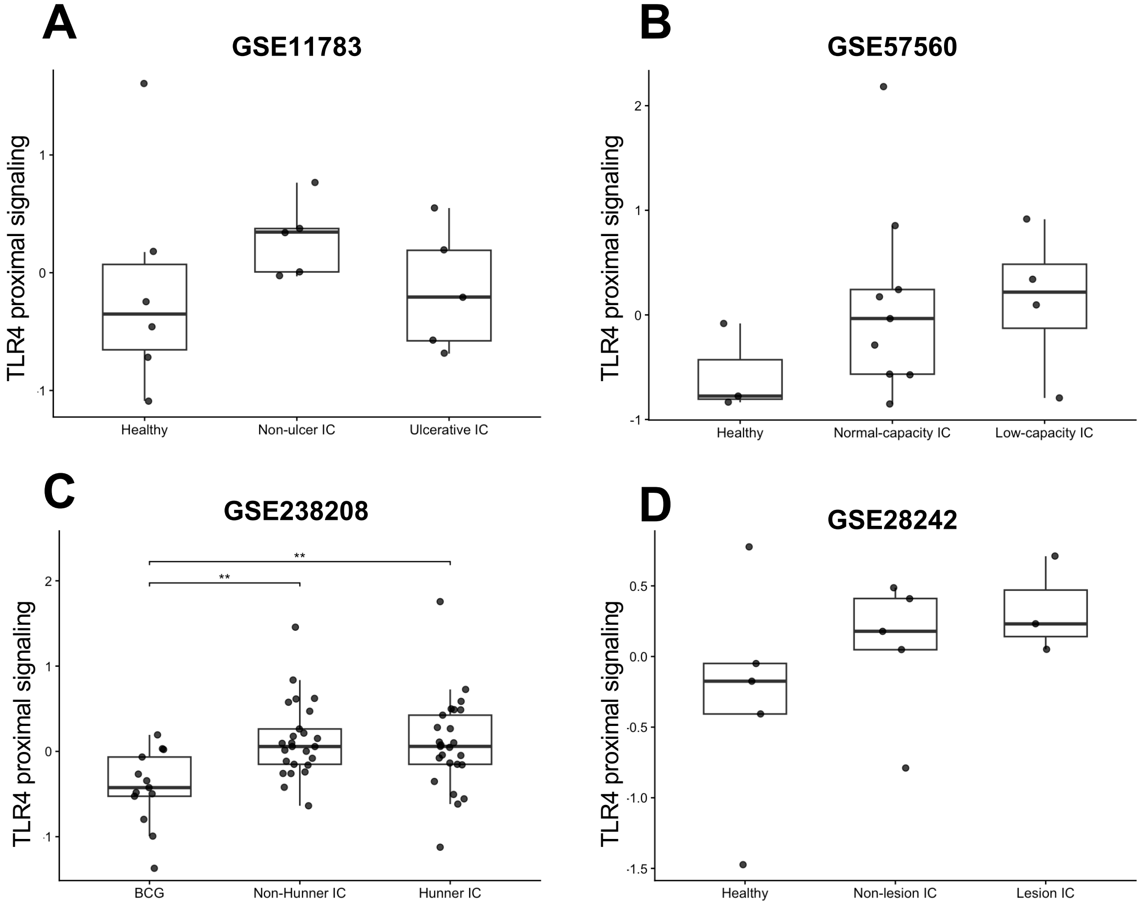

### Supplementary Figure 11.

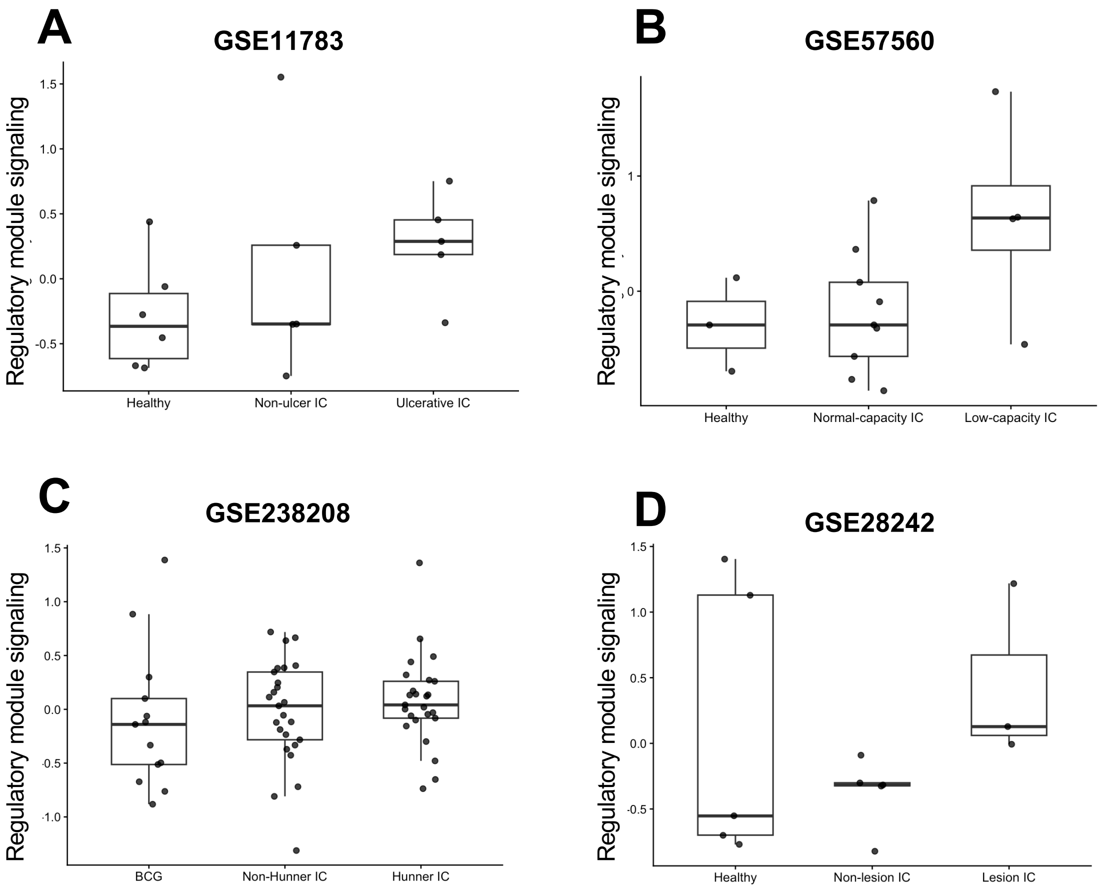

### Supplementary Figure 12.

A

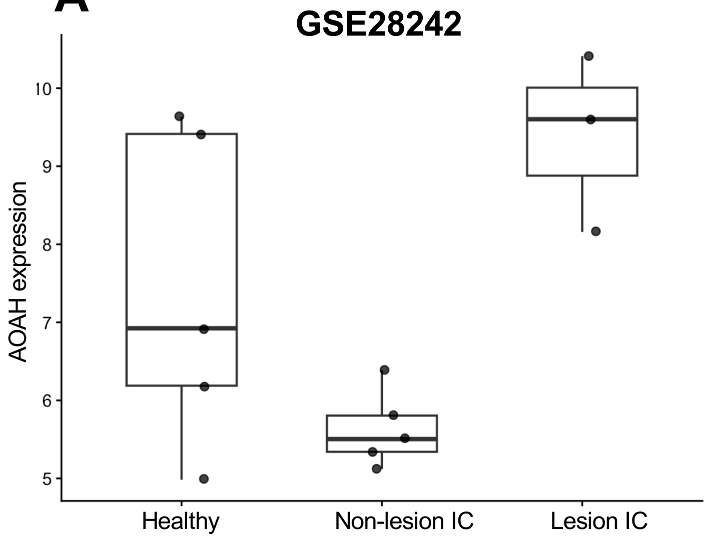

### Supplementary Figure 13.

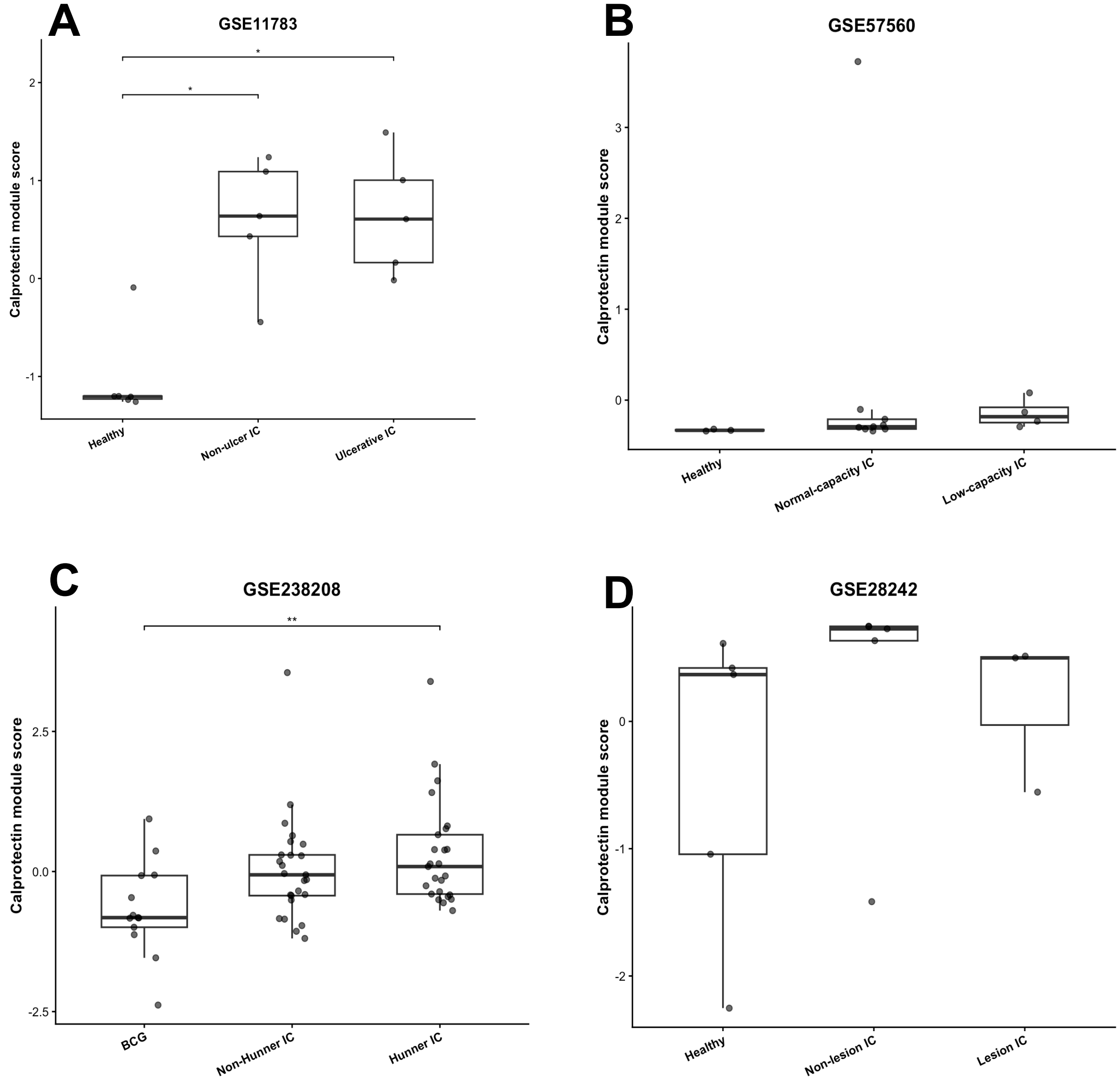

### Supplementary Figure 14.

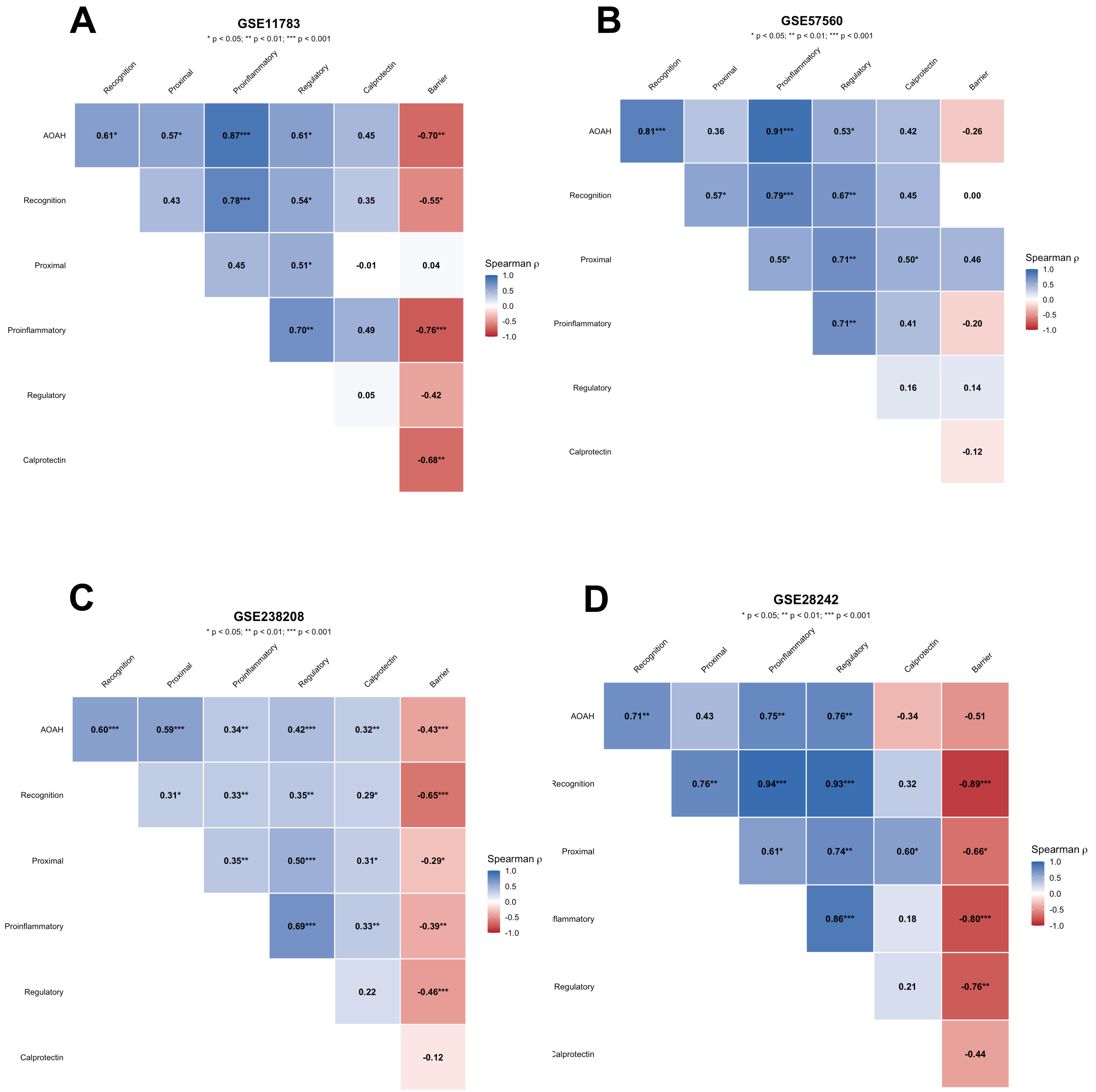

### Supplementary Figure 15.

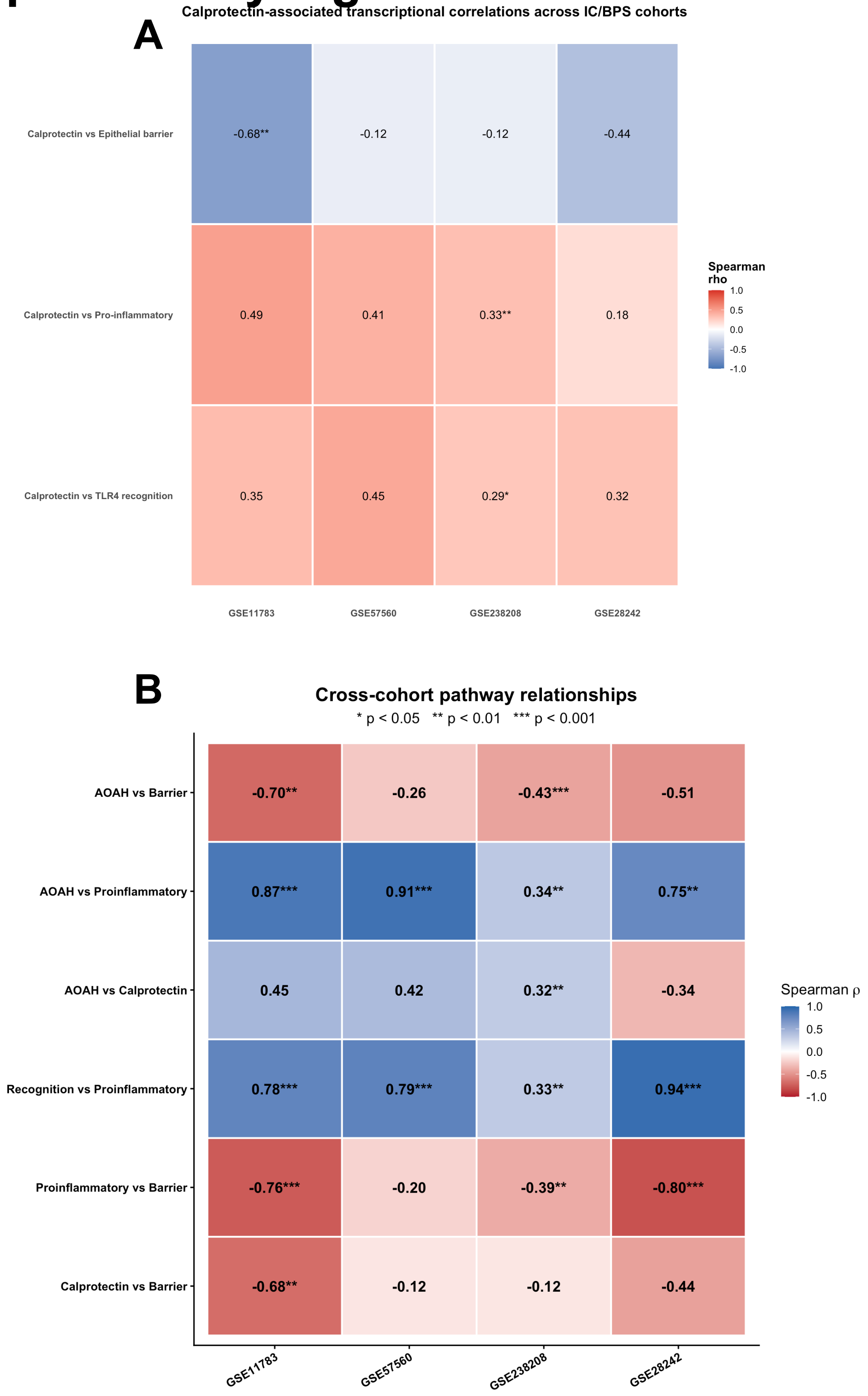

### Supplementary Figure 16.

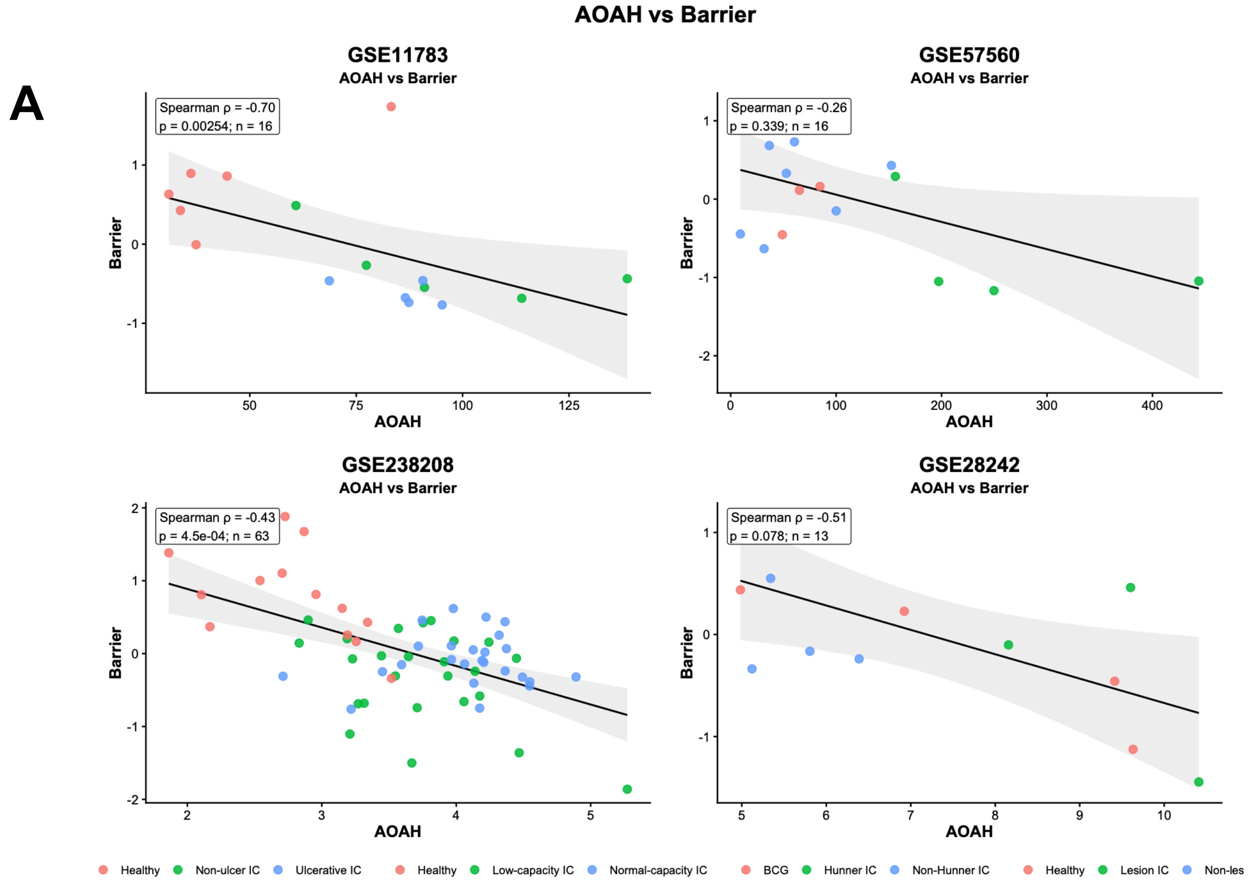


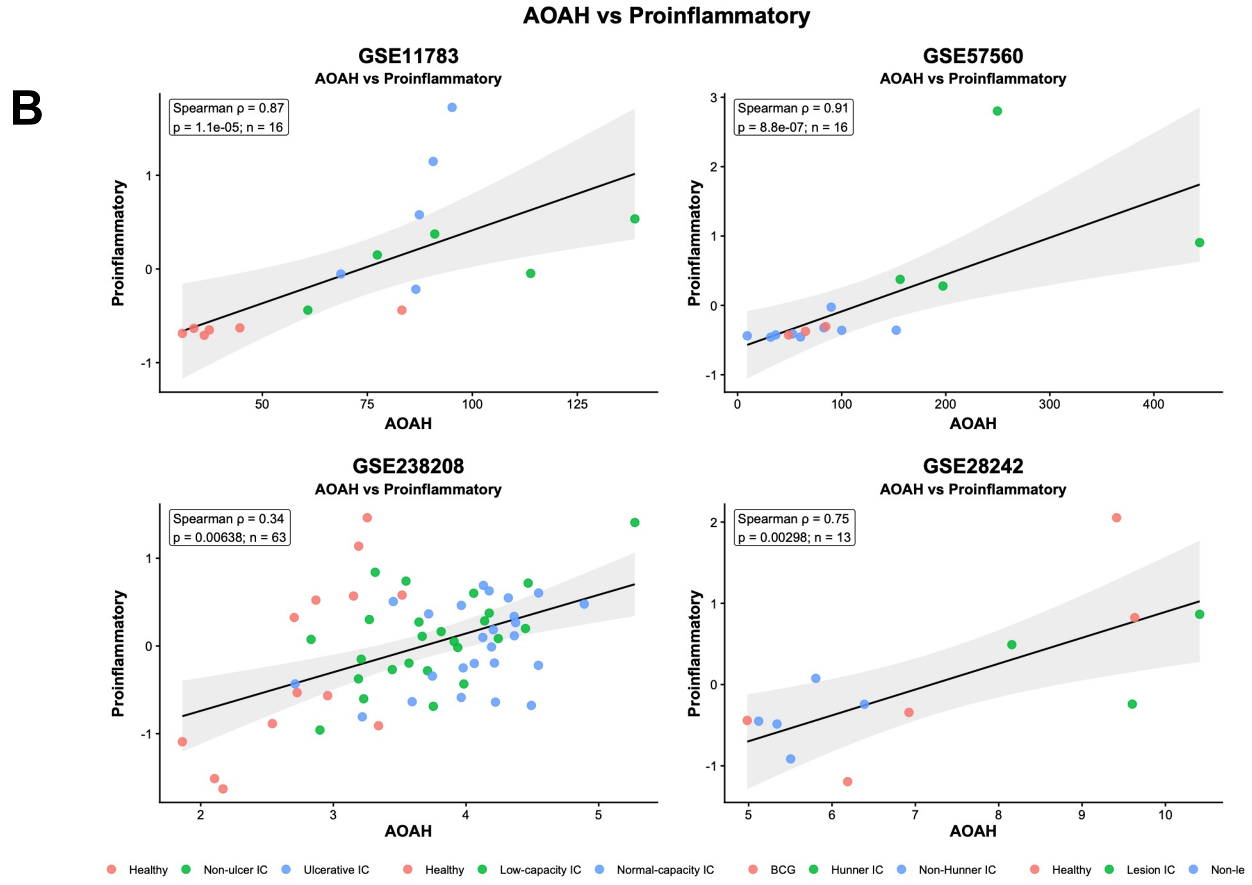

### Supplementary Figure 16.

C

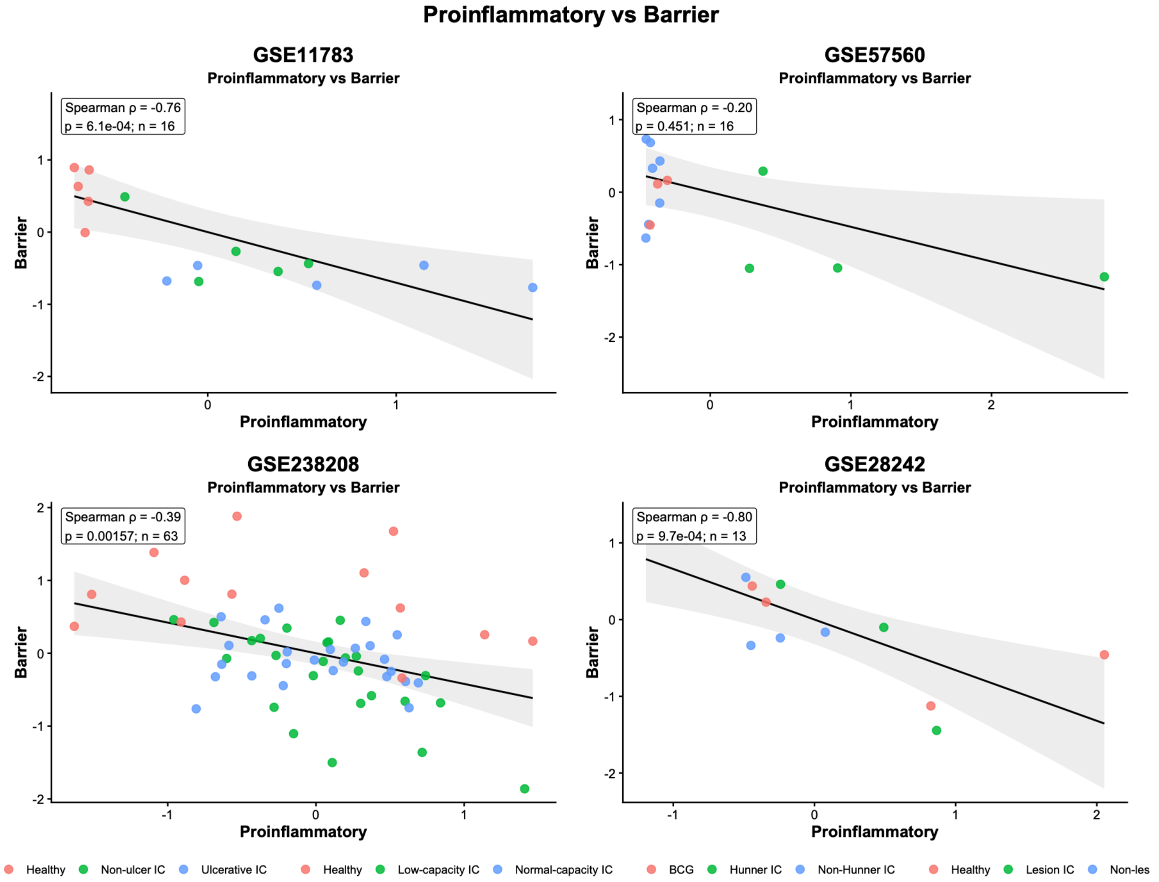


D

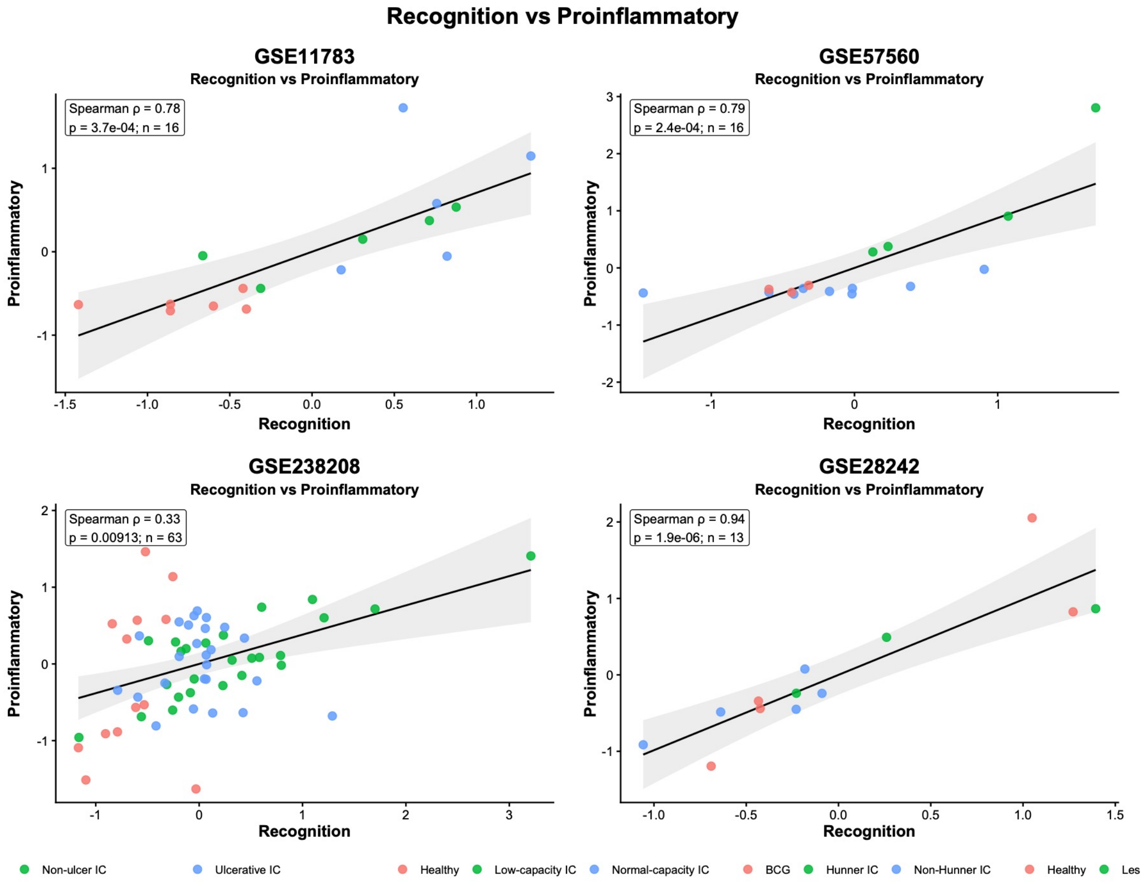

### Supplementary Figure 16.

E

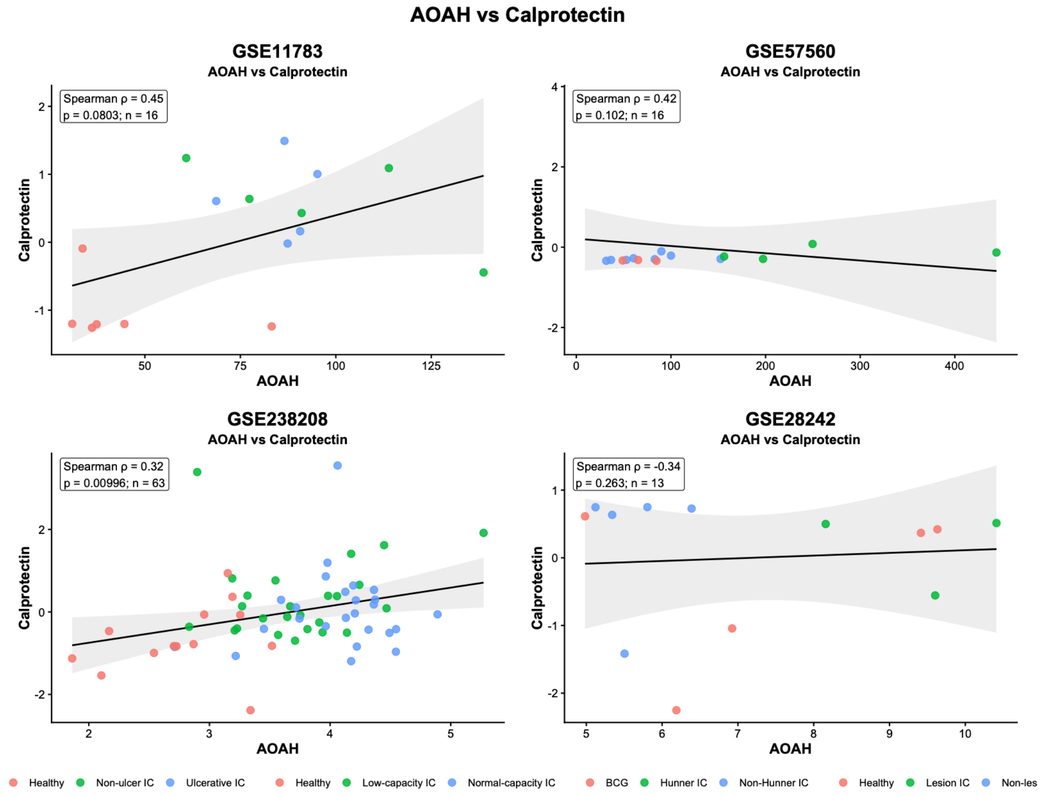


F

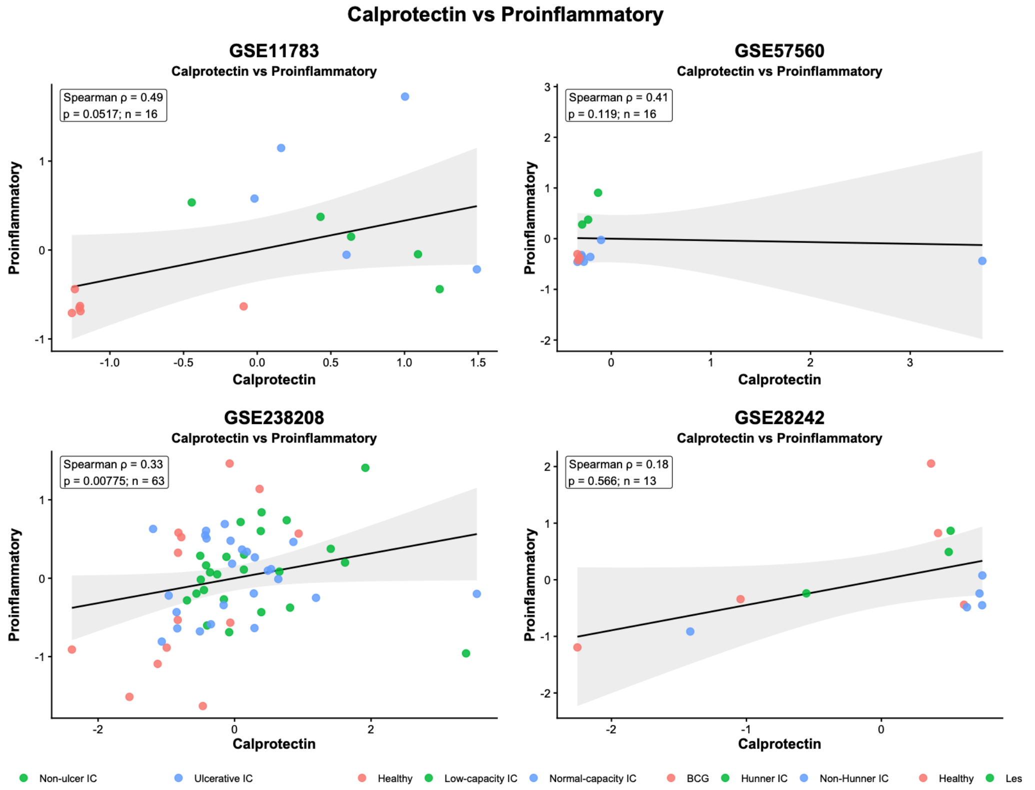

### Supplementary Figure 16.

G

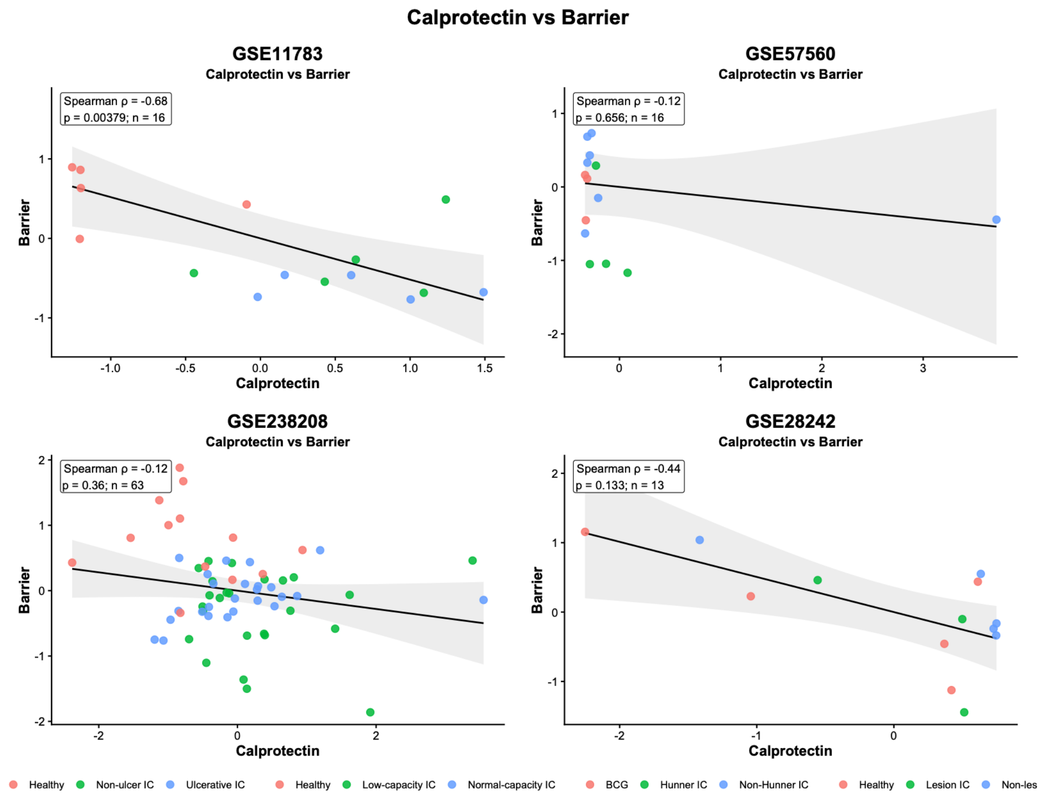

### Supplementary Figure 17.

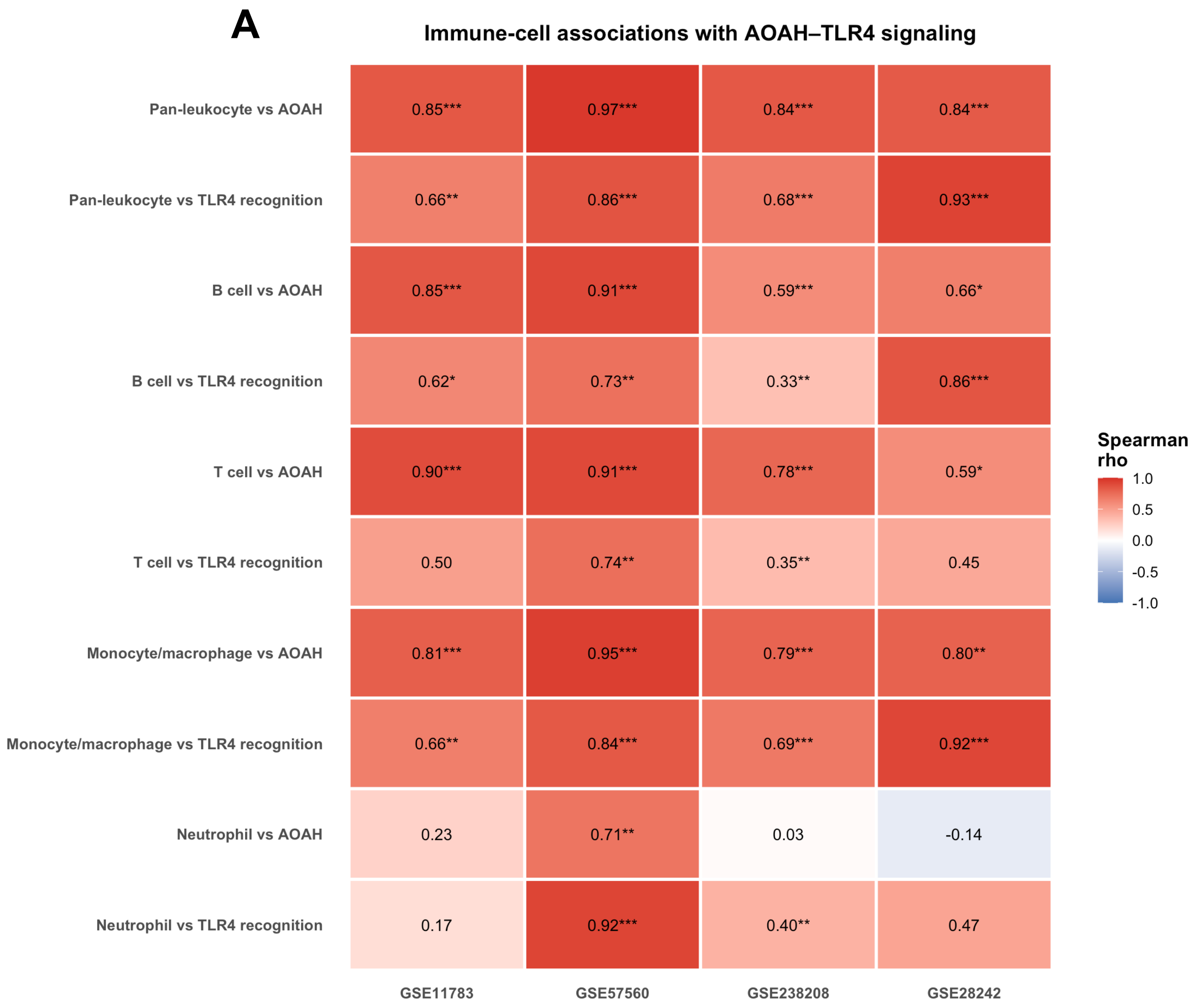

### Supplementary Figure 18.

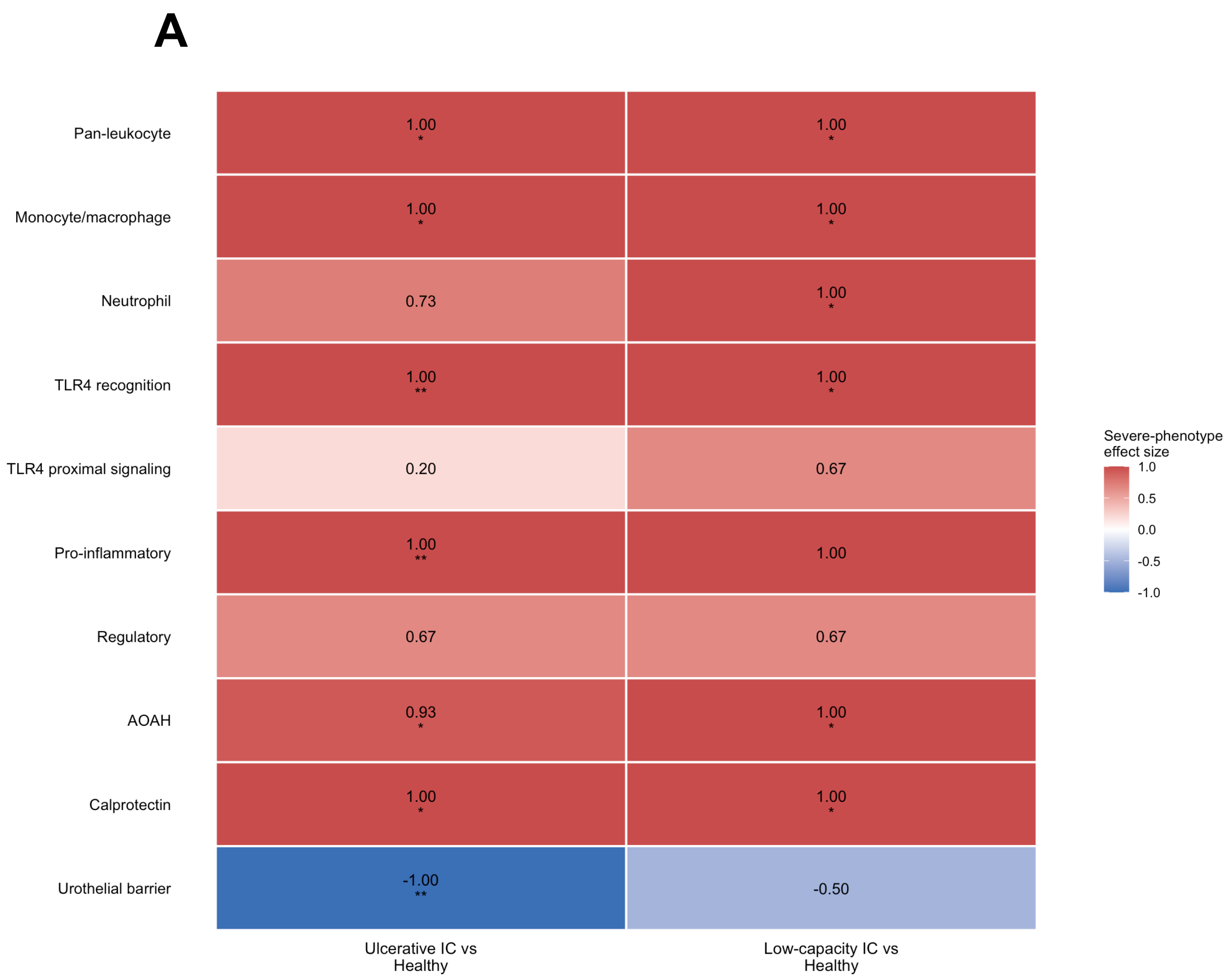

### Supplementary Figure 19.

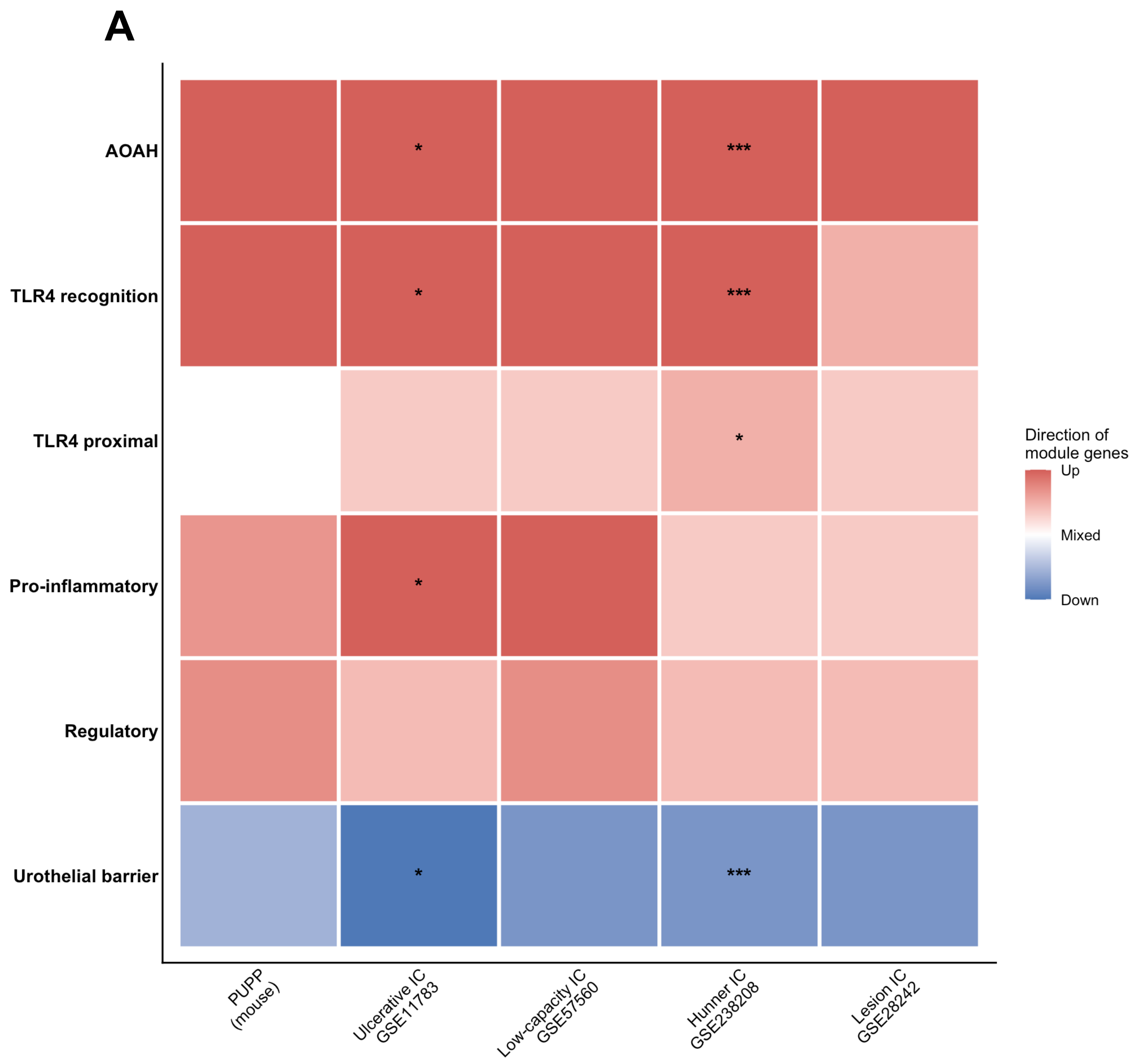
